# Unravelling Paralog-Specific Notch Signaling through Ternary Complex Stability and Transcriptional Activation Measurements Using Chimeric Receptors

**DOI:** 10.1101/2023.09.15.557970

**Authors:** Kristen M. Ramsey, Doug Barrick

**Author notes:** Lead Contact: Doug Barrick.

## Abstract

Notch signaling is mediated by four paralogous receptors with conserved architectures and overlapping yet non-redundant functions. Notch signaling generates a transcriptional activation complex (NTC) wherein the N-terminal RAM and ankyrin repeat (ANK) regions of the Notch intracellular domain (NICD) bind transcription factor CSL and recruit co-activator Mastermind-like (MAML). To better understand paralog-specific differences in Notch signaling, we analyzed the thermodynamics of binary and ternary NTCs for all four Notch paralogues and chimeric constructs. We find that RAMANK:CSL stability (ΔG_RA_) is primarily determined by the RAM region while MAML binding to preformed RAMANK:CSL complexes (ΔG_MAML_) is largely determined by the ANK region. We determined transcriptional activation data for the paralogous and chimeric NICDs and analyzed the data with an independent multiplicative model. This analysis shows ternary complex stability (ΔG_TC_, where ΔG_TC_= ΔG_RA_+ΔG_MAML_) correlates well with transcriptional activations and provides insights into contributions of RAM, ANK and the C-terminal regions to Notch signaling.

## Introduction

The Notch pathway is a highly conserved juxtacrine signaling pathway in metazoans used for communication between adjacent cells to regulate cellular differentiation in a context-dependent manner^1^. Its ability to signal between cells within a particular niche makes it a critical player in development in almost all animal organs and tissues^2,3^. Due to its key role in cell fate determination, dysregulation of Notch signaling by mutation and/or altered expression leads to a variety of disease states with onsets varying from intrauterine gestation to adulthood^4^. Defective Notch signaling during embryogenesis leads to significant congenital abnormalities^5^ while adult-onset conditions arising from altered Notch signaling include catastrophic vascular pathologies^6^ and a wide-range of cancers^7^, where Notch can act as an oncogene or tumor suppressor depending on the type of cancer^8–10^.

The Notch receptor is expressed as a 300 kDa, single-pass type I transmembrane protein composed of a ligand-binding extracellular domain (NECD), a short transmembrane helix, and an intracellular domain (NICD)^11^. Canonical Notch signaling is initiated by binding of the of the NECD to a Notch ligand on the surface of an adjacent cell, which triggers a series of proteolytic cleavages leading to endocytosis of NECD by the signal-transmitting cell and liberation of NICD from the cytosolic side of the plasma membrane in the signal-receiving cell^12^. Once released from the membrane, cleaved NICD translocates to the nucleus where it interacts with the DNA-bound transcription factor CSL, displacing co-repressors and recruiting the co-activator Mastermind-like (MAML) to form the ternary Notch transcriptional activation complex (NTC) which recruits transcriptional machinery to the promoters of Notch target genes (Figure 1A)^13^.

**Figure 1.**
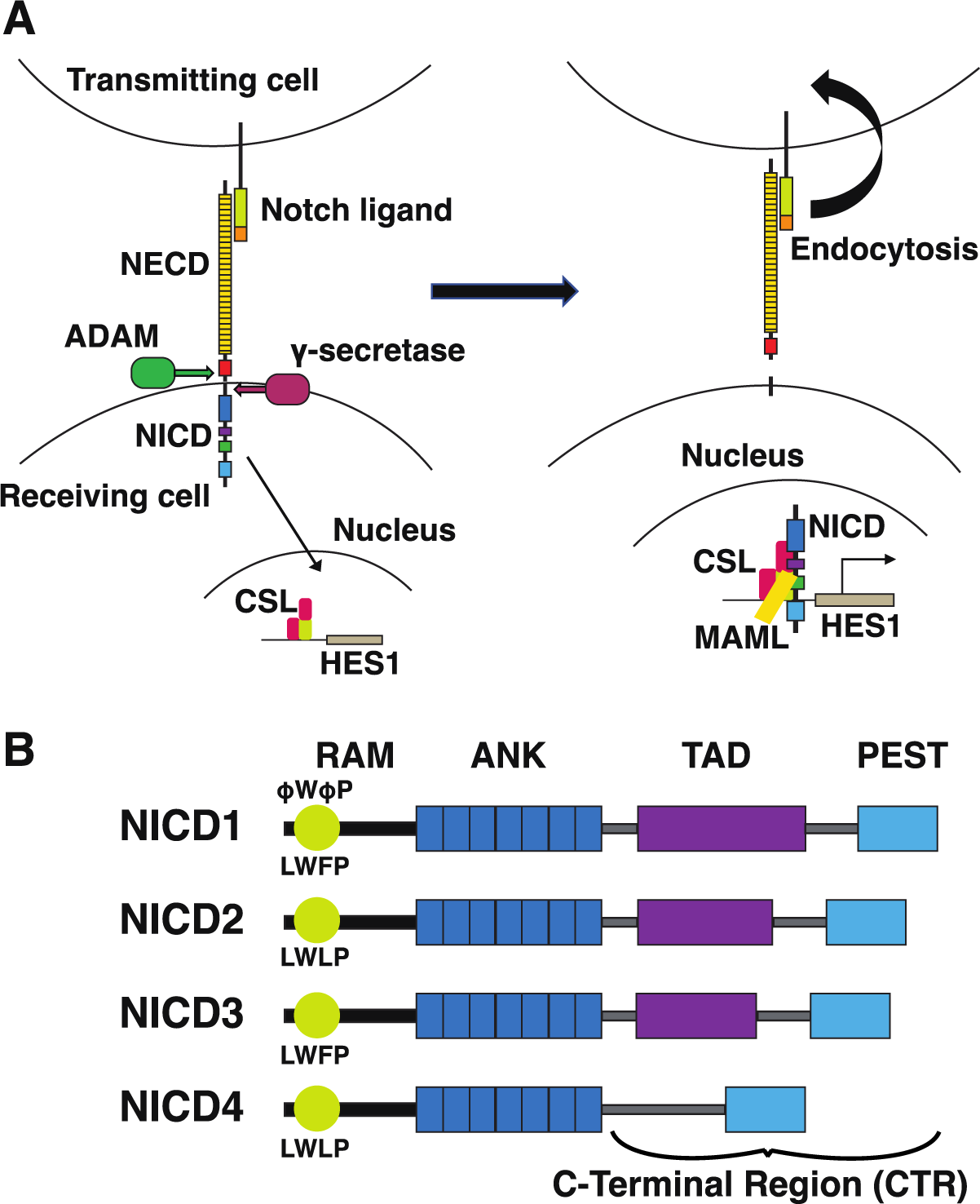
Canonical Notch transcriptional activation pathway and human NICD paralog domain organization. A) In canonical Notch signaling, the membrane bound Notch receptor interacts with a membrane-bound Notch ligand on an adjacent cell which triggers extracellular and intracellular proteolysis. The liberated Notch intracellular domain (NICD) then enters the nucleus, where it activates transcription through the transcription factor CSL and recruitment of co-activator Mastermind (MAML). B) Human NICD paralogs share a conserved architecture. A FWFP-containing N-terminal sequence within the RAM region binds to CSL and displacing co-repressors. The tandem ankyrin repeat domain (ANK) binds CSL and MAML. Notch1 and 2 contain C-terminal TAD regions, Notch 3 contains an unconventional TAD, and Notch4 lacks a C-terminal TAD. The C-terminal PEST sequence of all four Notch paralogues is responsible for degrading the NICD and terminating Notch signaling.

In humans, there are four Notch receptor paralogues (Notch1-4) which share the same canonical activation pathway^2^. Each of the four NICDs is organized in the same overall manner: at the N-terminus is a RAM (RBP-j-associated molecule) motif containing a conserved φWφP sequence (where φ stands for hydrophobic residue) critical for CSL binding, followed by a disordered region of 81 to 93 residues which links to seven tandem ankyrin repeats (ANK), and a C-terminal region (CTR) of variable length that includes a PEST degradation sequence (Figure 1B)^13–15^. The only large-scale differences between the paralogues lies in the region between the ANK domain and the PEST sequence where NICD1 contains a strong transcriptional activation domain (TAD) and NICD2 contains a weaker TAD^16^; NICD lacks a TAD in this region; likewise, NICD3 does not contain a conventional TAD but is proposed to interact with an unidentified zinc-finger transcription factor that contributes to the activation of specific Notch target genes, such as *Hes5*^17^.

Extensive studies of formation of the Notch1 NTC have shown that NICD1 forms a bivalent interaction with CSL through its RAM and ANK regions which bind at spatially distinct sites^18^. The highly conserved φWφP motif at the N-terminus of RAM binds with high affinity to the β-trefoil domain (BTD) of CSL^14^ while ANK binds to the remaining N- and C-terminal domains of CSL^14,15,18^. The N-terminal 80 residue region of MAML then binds to a cleft created between the NICD ANK and CSL, forming an extended α-helix that stabilizes the ANK:CSL interaction^15,18,19^.

In contrast to the well-characterized Notch1 NTC, there has been surprisingly little *in vitro* quantitative characterizaton of the NTCs of Notch2-4. Using cell culture and transgenic animals, several studies have shown that each paralogue activates different promoter architectures, and the origins of these differences have been mapped by substituting domains from one paralogue to another^17^. For example. studies in COS-7 cells have shown that weak activation of *Hes* promoters by Notch3 can be increased by replacing the ANK domain with that from NICD1 (a strong activator)^20^. Likewise, studies of T-cell acute lymphoblastic leukemia (T-ALL) induction show that the leukemic potential of NICD1 is eliminated when ARs 2-7 are substituted with those from NICD4^21^. These observations suggest that NICD paralogue specificity is controlled in a modular fashion.

Here we quantitatively and systematically define the role of individual domains in paralogue-specific NTC formation and transcriptional activation. First we used ITC to determine the thermodynamics of formation of the binary RAMANK:CSL complex, and we determined the thermodynamics of binding MAML to preformed RAMANK:CSL complexes to form ternary RAMANK:CSL:MAML complexes for the four Notch paralogues. Together, these two binding reactions give the complete thermodynamics of ternary NTCs from monomeric components. We show that each paralogue binds to CSL and recruits MAML with a unique thermodynamic profile, forming binary and ternary complexes of varying levels of stability. Next, by creating chimeric RAMANK constructs we were able to demonstrate that the paralogue-specific differences in binary complex assembly energetics are primarily determined by the identity of the RAM region whereas differences in MAML binding to preformed RAMANK:CSL complexes are primarily determined by the identity of the ANK domain. Finally, we determined the transcriptional activation potential of each NICD paralogue and NICD chimeras in which we shuffled the RAM regions, ANK domains, and C-terminal (TAD- and PEST-containing) regions (CTRs) of the four paralogues. To quantify the transcriptional activation potential of the three regions (RAM, ANK, CTR) and the degree to which these three regions contribute additively, we developed a model to globally fit the native and chimeric transcriptional activation strengths. This analysis supports and provides a quantitative basis for previous work on paralogue-specific domains and their effects on signaling.

## Results

### Binding thermodynamics of Notch1-4 paralogues and chimeric RAMANK constructs to CSL

To quantify the contributions of paralogue-specific RAM and ANK interactions to NTC stabilities, we determined the free energies and enthalpies of binding of each paralog’s RAMANK to CSL using isothermal titration calorimetry (ITC). The four paralogous RAMANK constructs were titrated into CSL to form binary complexes, and were fitted with a heterodimer scheme:

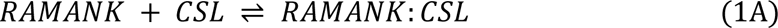

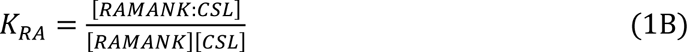

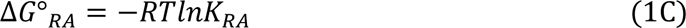

The heterodimer fitting equation includes an adjustable incompetent fraction of RAMANK (Figure 2A and Table 1). Titration data are well-fitted by the model, and fitted incompetent fractions of RAMANK were low (0.02 to 0.10), demonstrating that recombinant proteins are monomeric, well-folded, and active. The four paralogous RAMANK constructs form binary complexes with CSL with similar affinities (Table 1), with Notch1 having the lowest (most favorable) binding free energy (ΔG°_RA_ = −10.8 kcal mol^−1^, nearly identical to the value reported by Sherry et al^22^) and Notch3 RAMANK having the highest (least favorable) binding free energy (ΔG°_RA_ = −9.6 kcal mol^−1^). Though differences among paralogues are modest (1.1 kcal mol^−1^) they are statistically significant.

**Figure 2.**
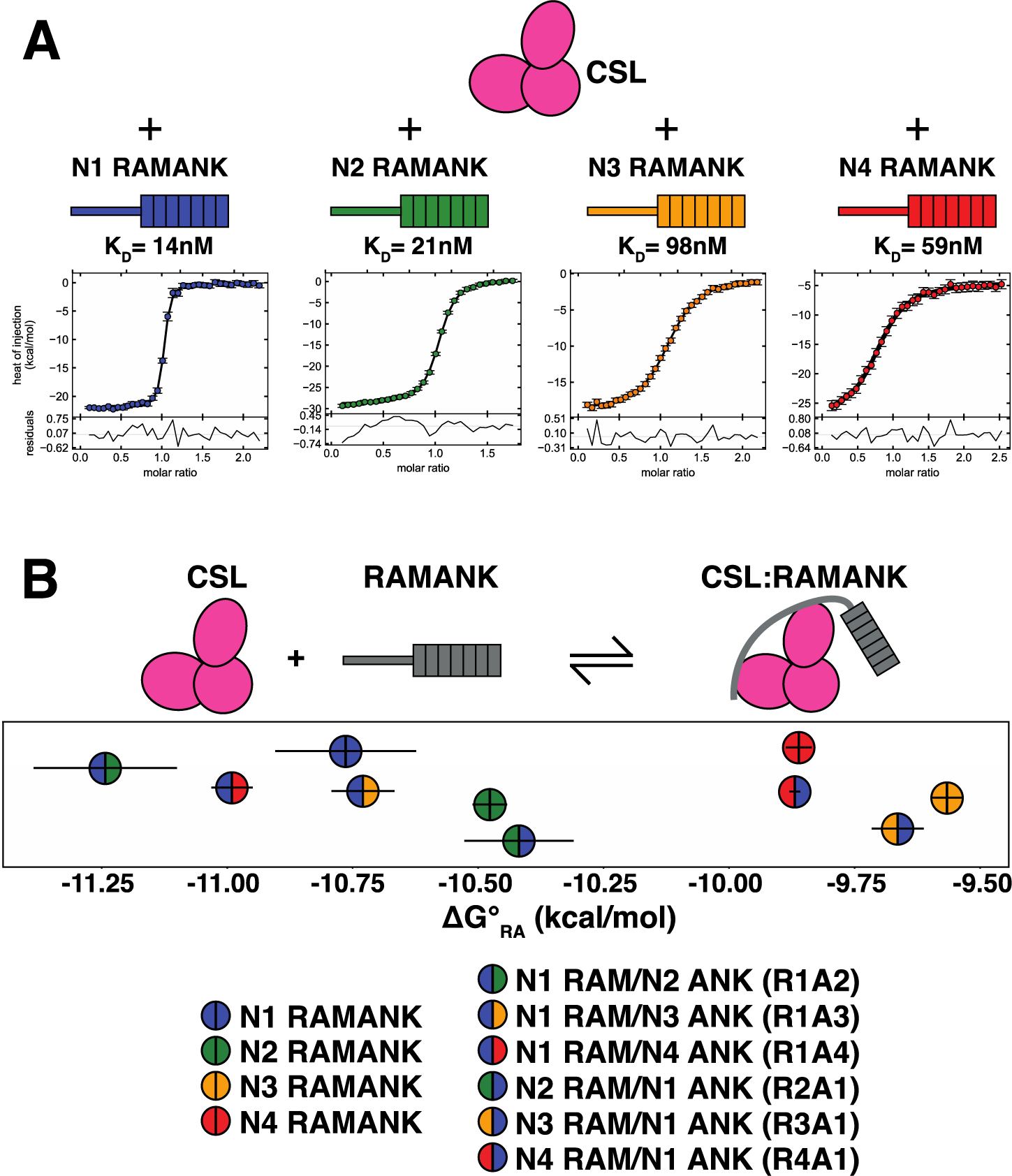
Isothermal titration calorimetry of cognate and chimeric RAMANK constructs binding to CSL. A) Each paralogue’s RAMANK domain was titrated into an ITC cell containing CSL. Resulting isotherms were fit using a single-site binding equation to determine thermodynamics of association. B) ΔG°_RA_ values for binding of RAMANK to CSL for the four Notch paralogues along with values for chimeric RAMANK constructs (Figure S1). Error bars report the standard error of the mean from at least three independent experiments. In general, chimeric RAMANK constructs bind to CSL with the free energy of the RAM domain’s paralogue. Throughout, Notch1, 2, 3, and 4 are colored blue, green, orange, and red, respectively. Thermograms for chimeric RAMANK constructs binding to CSL are shown in Figure S1.

**Table 1.** Thermodynamics of RAMANK paralogues and chimeras binding to CSL.

| | $K_D$ (nM)<sup>a</sup> | $\Delta G^{\circ}_{RA}$ (kcal mol<sup>-1</sup>)<sup>a</sup> | $\Delta H_{RA}$ (kcal mol<sup>-1</sup>)<sup>a</sup> | $T\Delta S_{RA}$ (kcal mol<sup>-1</sup>)<sup>a</sup> |
| --- | --- | --- | --- | --- |
| R1A1 | 14 ± 3 | -10.77 ± 0.11 | -20.56 ± 0.37 | -9.80 ± 0.28 |
| R2A2 | 21 ± 1 | -10.48 ± 0.03 | -26.54 ± 1.7 | -16.06 ± 1.7 |
| R3A3 | 98 ± 3 | -9.56 ± 0.03 | -21.60 ± 0.83 | -12.04 ± 0.81 |
| R4A4 | 59 ± 2 | -9.86 ± 0.03 | -22.80 ± 0.12 | -12.94 ± 0.14 |
| R1A2 | 6 ± 1.2 | -11.24 ± 0.14 | -30.17 ± 2.7 | -18.92 ± 2.8 |
| R1A3 | 14 ± 1.5 | -10.73 ± 0.07 | -30.24 ± 1.0 | -19.69 ± 0.98 |
| R1A4 | 9 ± 0.7 | -10.99 ± 0.04 | -28.00 ± 1.1 | -17.01 ± 1.1 |
| R2A1 | 24 ± 4 | -10.42 ± 0.11 | -20.08 ± 0.73 | -9.66 ± 0.76 |
| R3A1 | 83 ± 7 | -9.66 ± 0.05 | -19.71 ± 1.0 | -10.05 ± 1.0 |
| R4A1 | 59 ± 1.2 | -9.87 ± 0.01 | -20.62 ± 1.3 | -10.76 ± 1.2 |

<sup>a</sup>Uncertainties are standard errors of the mean from at least three independent measurements.

Because the interaction of the Notch1 ANK domain with CSL has been shown to require MAML^14^, differences in binary affinities among the four paralogous RAMANK constructs likely reflect differences in interaction of the RAM region with CSL. To test this, we created the chimeric RAMANK constructs that combine the Notch1 RAM with ANK domains from Notch2, 3, and 4 as well as constructs that combine the Notch2, Notch3, and Notch4 RAM regions with the ANK domain from Notch1. We used ITC to determine the free energies of chimeric RAMANK:CSL binary complexes (Figure 2B, Figure S1, and Table 1). All three chimeras containing the Notch1 RAM region have binary free energies similar to the Notch1 RAMANK paralogue, and are the lowest (most stable) of the chimeras (Figure 2B); likewise, chimeras with Notch2, 3, and 4 RAM regions have binary free energies quite similar to those of Notch2, 3, and 4 RAMANK constructs, respectively, with the Notch2 chimera binding with an intermediate free energy, and Notch4 and particularly Notch3 binding with high (least stable) free energies. These results support the hypothesis that binary affinities of four paralogous RAMANK constructs with CSL are determined by the RAM region.

### Binding thermodynamics of MAML to paralogous and chimeric Notch1-4 RAMANK: CSL complexes

Since the free energy of formation of the RAMANK:CSL binary complex is determined largely by the RAM region, and since the isolated RAM region of Notch1 binds to CSL with a nearly identical free energy to that of Notch1 RAMANK^22,23^, the ANK domain is likely dissociated from its binding site on CSL in the RAMANK binary complex. To the extent that this is true, pre-bound RAMANK:CSL complexes provide a simple route to interrogating the thermodynamics of ANK binding to its site on CSL: titrating these binary complexes with MAML should produce a ternary complex in which the tethered ANK domain recruits MAML to CSL, forming a ternary complex according to the scheme

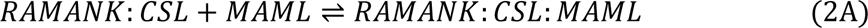

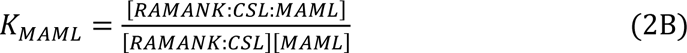

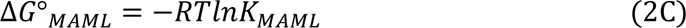

Because all four paralogous RAMANK constructs bind to CSL with affinities of 100 nM or below, pre-binding CSL at a concentration of 2 μM with a ten-fold excess of RAMANK will lead to greater than 99% saturation at the RAM binding site on CSL. As the ANK domain is tethered *in cis* in binary RAMANK:CSL complexes, MAML titrations of pre-bound RAMANK:CSL binary complexes should conform to a simple heterodimer binding scheme in which MAML forms a 1:1 complex with RAMANK:CSL. To the extent that the ANK domain is dissociated from its binding site on CSL in the RAMANK:CSL complex, the MAML binding free energy will report on the free energy of the ANK:CSL interface. In addition, ΔG°_MAML_ will also include contributions from the MAML:ANK and the MAML:CSL interfaces, although the latter contribution will be the same for all Notch paralogues and chimeras. In addition, ΔG°_MAML_ may be influenced by conformational changes within the RAM linker.

ITC titrations of RAMANK:CSL complexes with MAML are well-fitted by a heterodimer model for all four Notch paralogues. Fitted free energies are similar in magnitude to those of the binary complex (an average of about −11 kcal mol^−1^, Figure 3, Table 2). However, enthalpies are significantly larger in magnitude (more negative), reflecting a larger enthalpy-driven reaction opposed by a larger entropy penalty (Table 2). This enthalpy/entropy distribution may be the result of a disorder-order transition associated with MAML binding: crystal structures of the NTC have shown MAML to adopt an extended α-helix^14,15,18^, whereas secondary structure predictions indicate that MAML is disordered when free in solution^19^. In addition, this may involve ordering of the RAM linker upon MAML binding. As with binary RAMANK:CSL complexes, the Notch1 RAMANK:CSL complex has the most favorable MAML binding free energy (ΔG°_MAML_ = - 11.7 kcal mol^−1^). In contrast to the free energies of RAMANK:CSL formation, the MAML binding free energy to Notch3 RAMANK:CSL complex is more favorable than to the Notch2 and 4 complexes (ΔG°_MAML_ = −11.3 versus −10.25 and −10.85 kcal mol^−1^).

**Figure 3.**
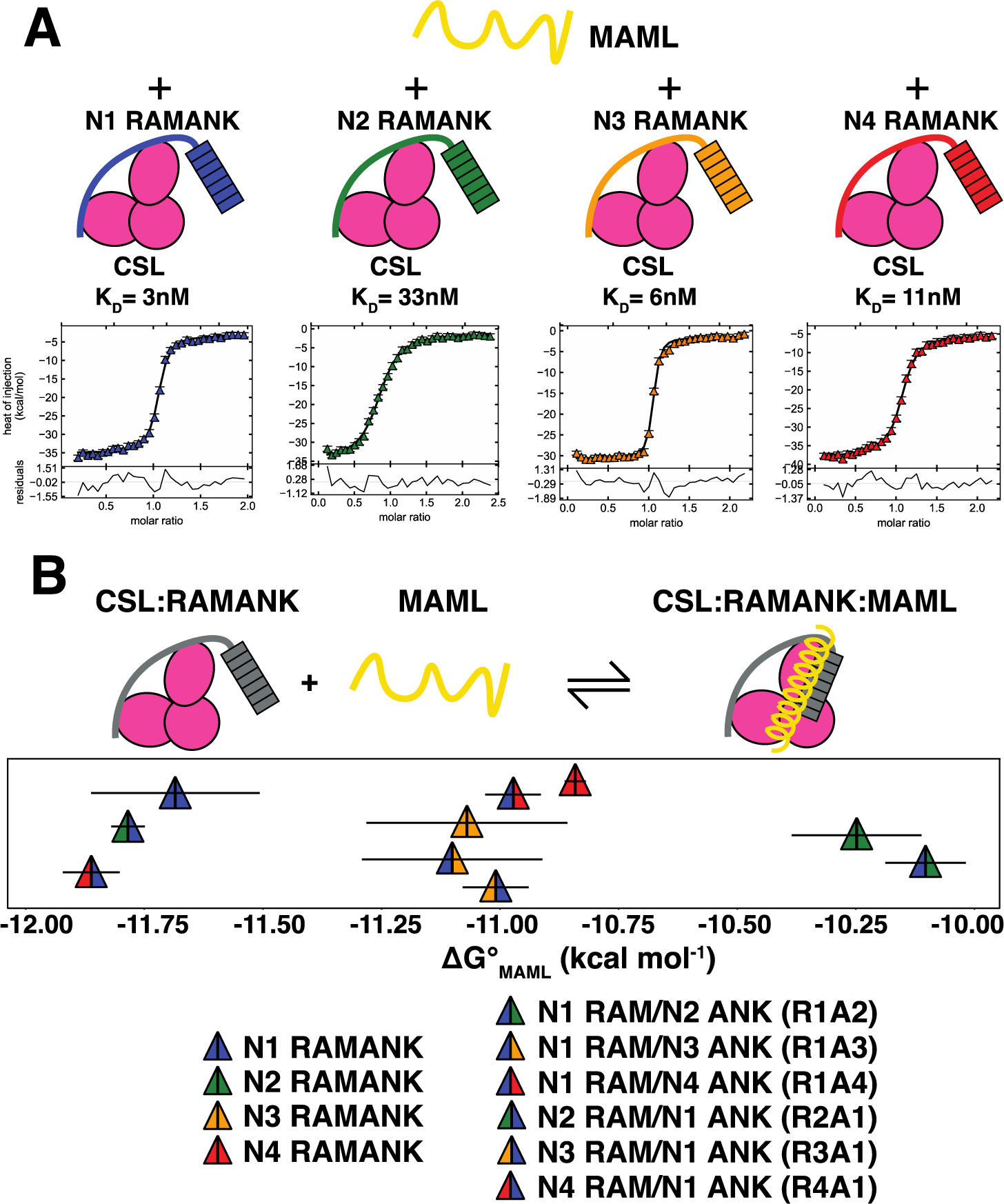
Isothermal titration calorimetry of MAML binding to pre-formed CSL:RAMANK paralogues and chimeras. A) MAML was titrated into an ITC cell containing CSL and a ten-fold molar excess of each RAMANK. Resulting isotherms were fit using a single-site binding equation to extract thermodynamic parameters. B) ΔG°_MAML_ values for binding of MAML to pre-formed CSL:RAMANK complexes for the four Notch paralogues along with values for chimeric RAMANK constructs. Error bars report the standard error of the mean from at least three independent experiments. With the exception of R3A1, chimeric RAMANK constructs in complex with CSL recruit MAML with thermodynamics resembling the ANK domain’s paralogue. Colors are as in Figure 2. Thermograms for MAML binding to chimeric RAMANK:CSL complexes are shown in Figure S2.

**Table 2.**
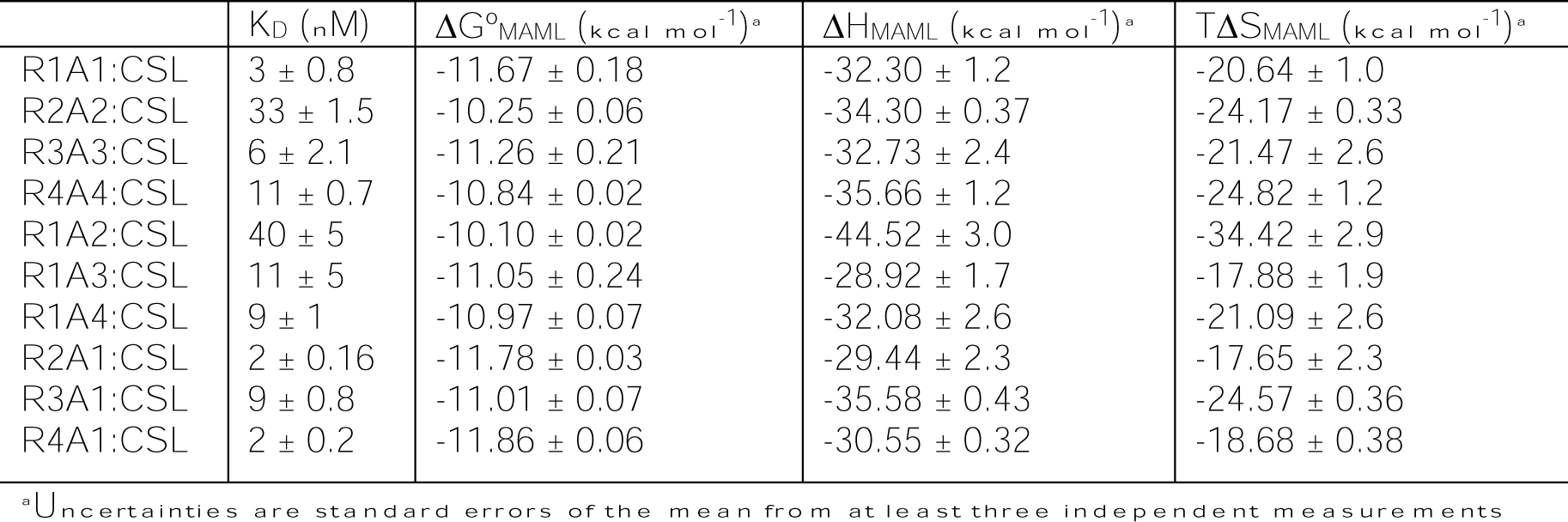
Thermodynamics of MAML recruitment to paralogous and chimeric RAMANK:CSL.

As described above, differences in free energies of MAML binding to paralogous RAMANK:CSL complexes are likely to originate in sequence differences among the ANK domains (rather than the RAM regions). To test this, we measured MAML binding to CSL with pre-bound RAMANK chimeras (Figure 3B, Figure S2, and Table 2). For the most part, chimeric RAMANK:CSL complexes have the same MAML binding free energies as Notch paralogues with the same ANK domain. For example, the chimera with RAM from Notch1 and ANK from Notch 2 (R1A2) has a high (unstable) MAML-binding free energy as does the Notch2 paralogue (ΔG°_MAML_ = −10.1 and −10.25 kcal mol^−1^, respectively). Likewise, R1A3 and R1A4 chimeras have intermediate ΔG°_MAML_ values, closely matching values for the Notch3 and Notch4 RAMANK constructs. Two of the chimeric constructs with the Notch1 ANK domain (R2A1 and R4A1) have low (stable) ΔG°_MAML_ values, matching that of the Notch1 RAMANK construct, although the third (R3A1) has an intermediate binding free energy. With the exception of the R3A1 chimera, these results support the hypothesis that MAML binding affinities to the four paralogous RAMANK:CSL complexes are determined by the ANK domain.

### Transcriptional activation by NICD paralogues and chimeras

To assess the potency of transcriptional activation by each paralogue, we performed dual luciferase assays in which we transfected HeLa cells with a plasmid encoding each paralogue’s NICD along with a plasmid containing the firefly luciferase gene downstream from 12 tandem CSL binding sites. Consistent with previously reported data^17^, transcriptional activation from NICD1 is twice that of NICD2, 3, and 4 (Figure 4). This ranking is consistent with findings from ITC that the RAM and ANK regions from Notch1 bind more tightly to CSL than the RAM and ANK regions from the other three paralogues.

**Figure 4.**
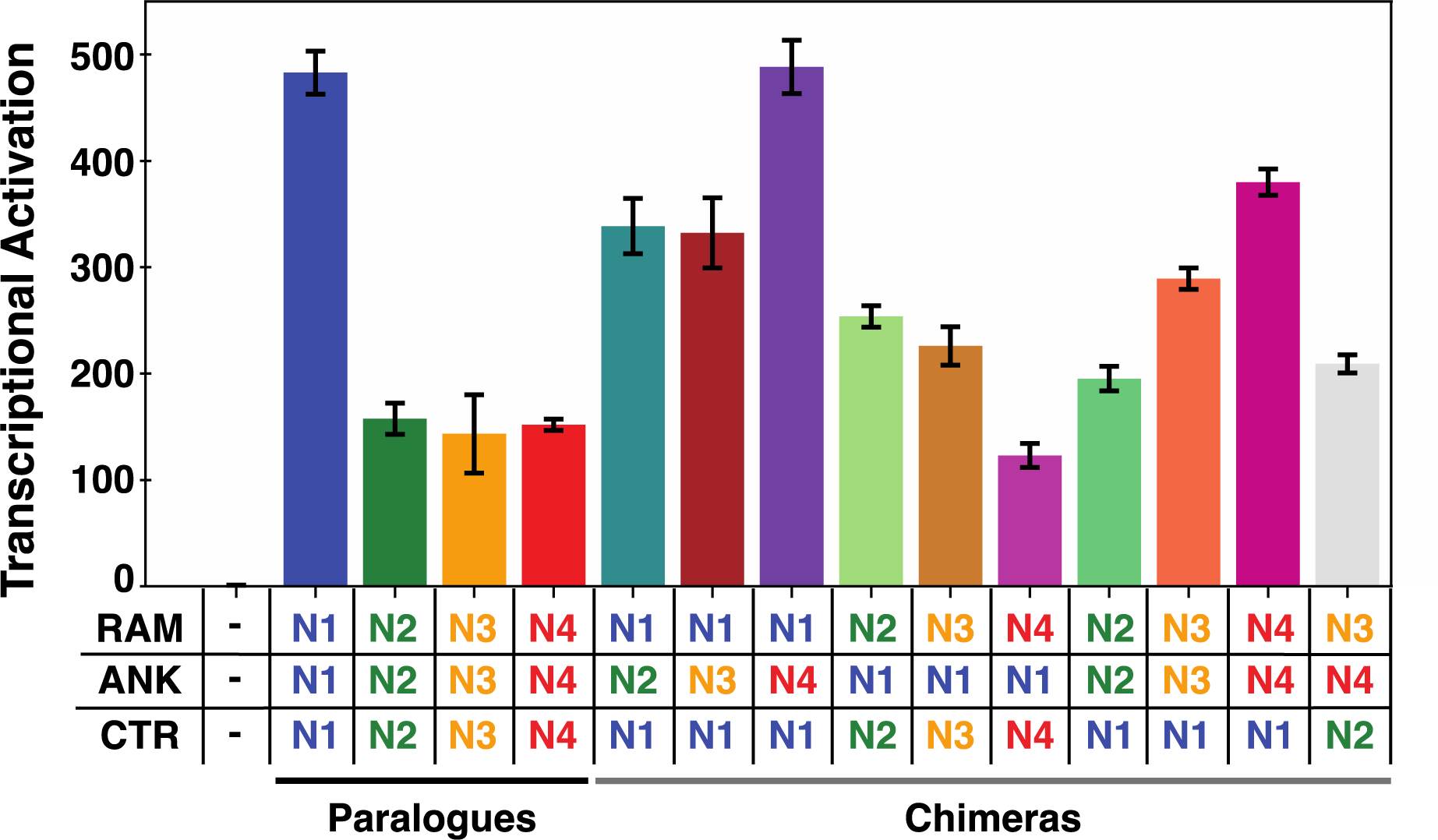
Transcriptional activation by the four NICD paralogues and their chimeras. NICD paralogues (left) and chimeras (right) display a range of activation levels on a synthetic Notch promotor driving luciferase transcription. Each bar represents the average of at least three independent transfections performed in triplicate. Error bars are the standard errors of the means. Expression levels were confirmed using Western blotting shown in Figure S5.

To resolve the contributions of the RAM and ANK regions of each paralogue to transcriptional activation, we created chimeric RAMANK NICD constructs for luciferase assays. Because the CTRs are known to contribute to transcriptional activation, and there are significant sequence differences among the four paralogues, our chimerization strategy for the NICDs included CTR variation. Along with the four paralogous NICDs, the chimeric NICDs show a considerable range of transcriptional activation. Likewise, all three regions (RAM, ANK, and CTR) appear to make contributions to transcriptional activation.

The contributions of the ANK domains to transcriptional activation can be seen most clearly by taking Notch1, which shows the highest level of transcriptional activation, as a reference. Swapping ANK domains from Notch2 and 3 (but not Notch4) into NICD1 decreases transcriptional activation. Likewise, swapping Notch1 ANK into NICD2 and 3 (but not NICD4) increases transcriptional activation. These comparisons indicate that the Notch1 ANK makes a stronger contribution to transcriptional activation than Notch2 and Notch3 ANKs, consistent with binding free energies of ANK and MAML to CSL (Figure 3), although the similar activation levels of Notch1 and Notch4 ANKs are at odds with the increased ΔG_MAML_ value (lower stability) for Notch4 ANK.

The contribution of the CTRs to transcriptional activation is illustrated in chimeras that swap the CTR of Notch1 into NICD2, 3, and 4 (Figure 4): all three of these chimeras (R2A2C1, R3A3C1, and R4A4C1) have increased transcriptional activation compared to NICD2, 3, and 4. (Figure 4). The increased activation by the Notch1 CTR is consistent with previous results that show the Notch1 C-terminal TAD to be strongly activating^16^. The finding that swapping the Notch1 CTR into NICD4 results in the largest increase in transcriptional activation is consistent with the lack of a TAD sequence in the CTR of Notch4.

Results from the chimeric NICD constructs demonstrate that the RAM, ANK, and CTR regions all contribute to the variability in transcriptional activation by the four Notch paralogues (Figure 4). To quantify the relative contributions of these three regions to transcriptional activation in a statistically rigorous way, we developed a model that could be fitted to transcriptional activation values of all fourteen constructs (four paralogues and ten chimeras). In addition to providing estimates for the contributions of individual regions to transcriptional activation, the quality of the fit provides information on whether the structure of the model adequately captures the variation in the data.

The simplest type of model that can be used to analyze the data is one in which the contributions of individual regions are independent of their context. In such a model, contributions of paralogous regions to transcriptional activation can either be additive, i.e.,

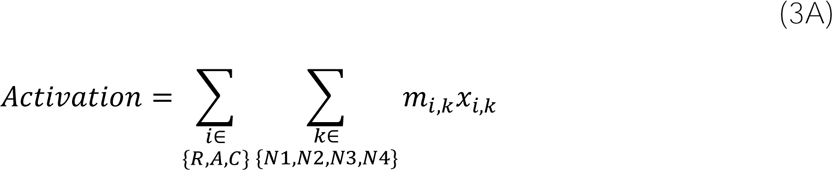

or multiplicative, i.e.,

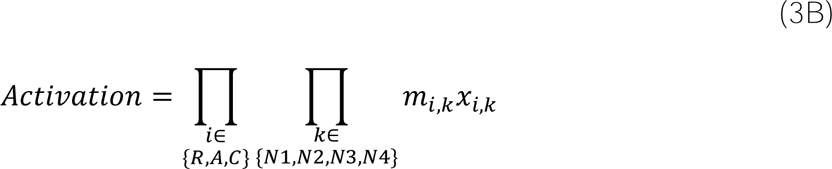

where *m_i,k_* values are unknown parameters proportional to the contribution of the *i*^th^ region from the *k*^th^ paralogue to activation, and *x_i,k_* are binary variables that have values of one if a construct includes region *i* from paralogue *k*, and zero otherwise. For example, the transcriptional activiation of chimera R1A4C2 would be given by the additive model as

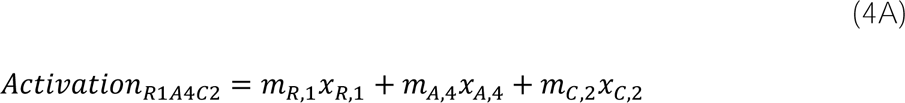

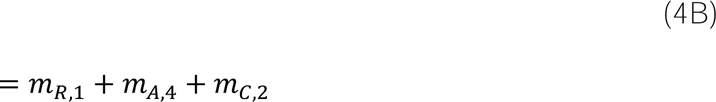

and by the multiplicative model as

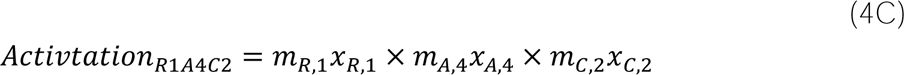

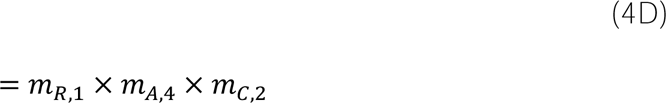

Transcriptional activation should be proportional to the extent of Notch ternary complex formation at the luciferase promoters, which at subsaturating NICD levels should be proportional to the overall binding constant for ternary complex assembly. This overall binding constant should in turn be proportional to the product of the binding constants for the RAM and ANK(+MAML) regions. Moreover, the amount of ternary complex will be proportional to the amount of nuclear NICD, which is determined, at least in part, by the C-terminal NLSs. Finally, the downstream activation afforded by the C-terminal TADs will be proportional to the extent of ternary complex formation, and thus to the strength of RAM and ANK(+MAML) interactions. Because all of these synergies would be captured by a multiplicative model, we analyzed the transcriptional activation values using this model. By taking the logarithms of the transcriptional activation data, the multiplicative model becomes linear. For example, the equation relating the activation of R1A4C2 to its RAM, ANK, and CTRs (equation 4D) becomes

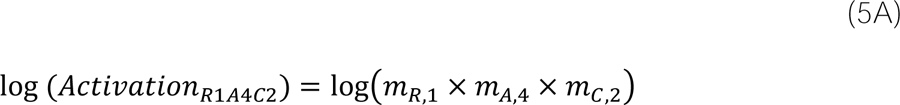

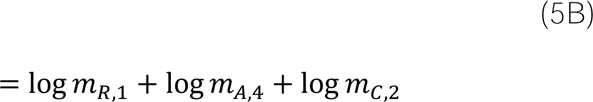

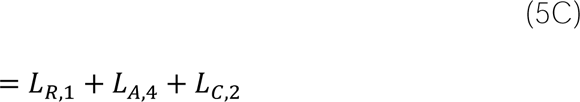

where the *L_i,k_* parameters are directly fitted to the log of transcriptional activation values using simple linear regression.

Because each of our fourteen constructs all contain one and only one of each of the three regions, there are strong correlations among parameters for the three regions. As a result, our data do not span the entire space of the model^1^. This correlation is eliminated by representing one paralog, NICD3, using a single adjustable parameter and fitting the contributions of the other three paralogues’ RAMs, ANKs, and CTRs relative to NICD3. This reparameterization decreases the number of unknown parameters from 12 to 10, and leaving 4 degrees of freedom. We found that the same trends in the fitted parameters were obtained regardless of whether NICD2, NICD3, or NICD4 was represented with a single parameter; we chose NICD3 for the analysis presented here because it generates slightly lower parameter correlation than the other paralogues (Supplemental Figure 3). With this modified ten-parameter model, activation is given by

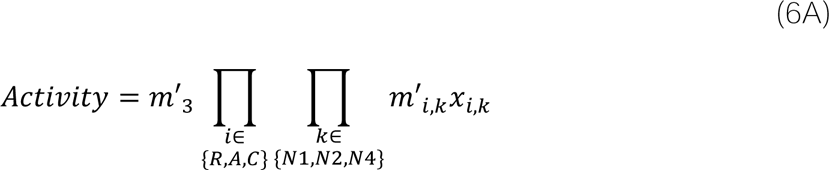

Where *m’_3_* gives the activity of NICD3, and the *m’_i,k_* parameters give contributions of the RAM, ANK, and CTRs of NICD1, 2, and 4 relative to NICD3. For example, the log of the activation of chimera R1A4C2 becomes

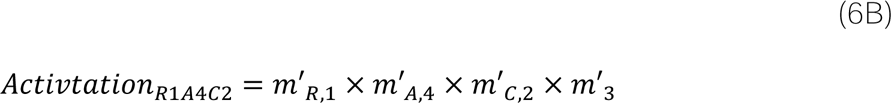

and in log space

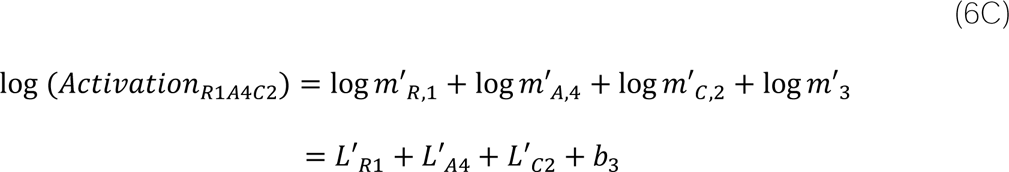

Because the model was fit to the log of activation values, fitted parameter values (e.g., *L*′_*R*1_) greater than zero correspond to an increase in transcriptional activation relative to NICD3 whereas values less than zero correspond to a decrease in relative activation. In terms of activation values, *m*′_*i,k*_ values greater than one indicate an increase in transcriptional activation relative to NICD3 whereas values less than one indicate a decrease in relative activation.

The *L’_i,k_* parameters generated from the fit using equation 6C (Figure 5A, Table 3) faithfully capture the measured transcriptional activation measured for each paralogue and chimeric construct (Figure 5B), with a Pearson correlation coefficient of 0.987 between predicted and measured activation levels. The activation parameters for the three regions of NICD1 are the largest among those from all three paralogues: values for the RAM and CTR of NICD1 exceed values from NICD2 and NICD4, and the positive coefficient values for all three regions of NICD1 indicate stronger activation than for the same regions from NICD3 (Figure 5A). The *L’_i,4_* parameters for all three regions of NICD4 display the largest range of contributions to activation with its CTR being the weakest contributor (both among the three Notch3 parameters and among the CTR paramaters for Notch1, 2, and 4) and its ANK domain contributing to activation at the same level as the NICD1 ANK.

**Figure 5.**
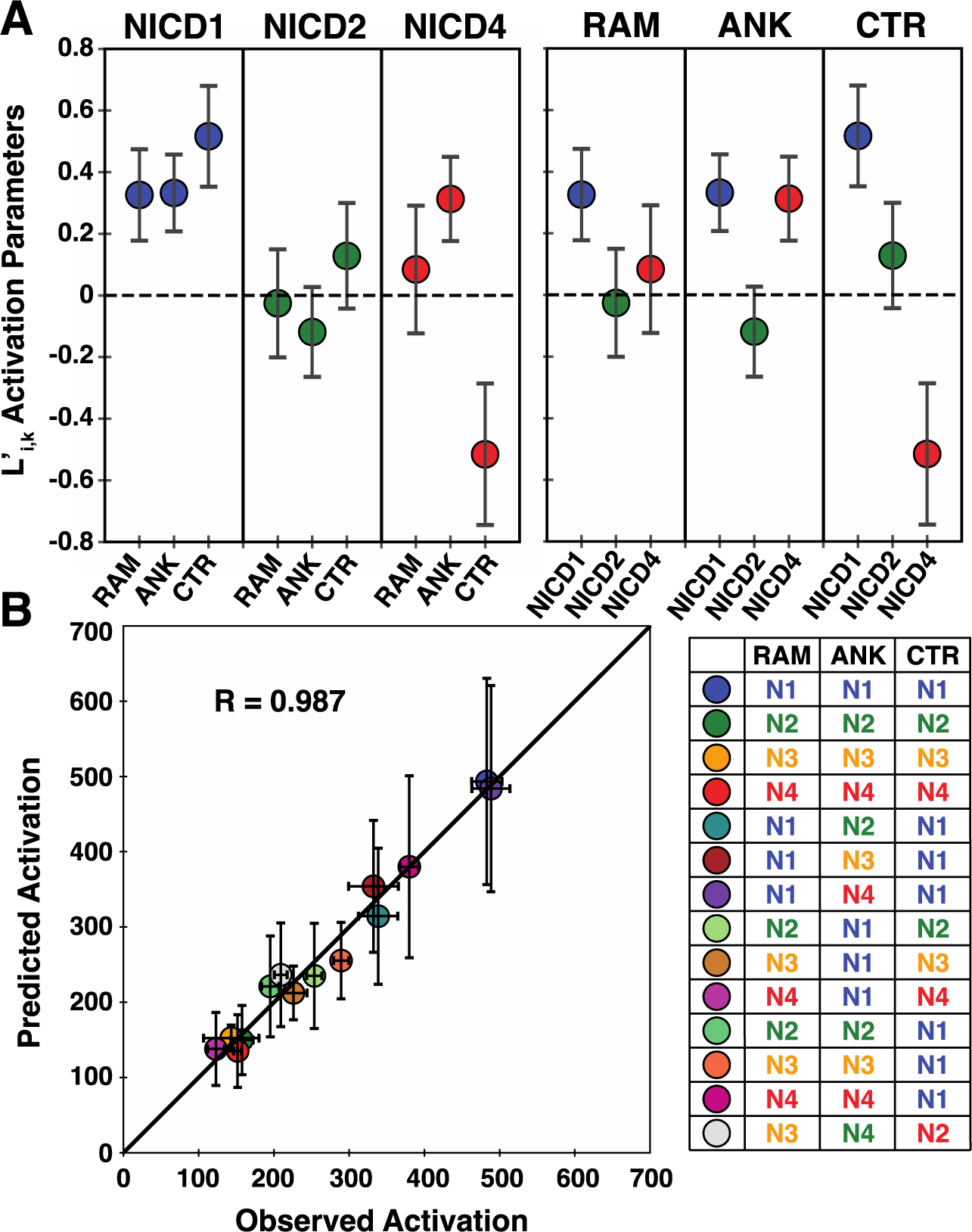
Contributions to transcriptional activation of the RAM, ANK, and C-terminal regions of the four Notch paralogues. A) The fitted *L’_i,k_* parameters (equation 6C) for the RAM, ANK, and CTR regions of NICD1, NICD2, and NICD4. Values are in log-space relative to NICD3. Values greater than zero increase transcriptional activation relative to NICD3, whereas values less than zero decrease relative activation. Parameters are grouped by paralog (left) and by region (right). Error bars represent the standard deviations calculated from 10,000 bootstraps of the fit. B) Correlation between the predicted transcriptional activation values of all the paralogs and chimeric NICDs constructs from the model with observed values. The R-value is a Pearson correlation coefficient. The color scheme is the same as in Figure 4. Error bars in the x dimension are +/- the standard error of the mean from the measured transcriptional activations as in Figure 4. Errors on predictions are calculated by propagating bootstrap errors using standard formulae^28^. See Figure S3 for covariance matrices for models fitting transcriptional activation data relative to NICD3, NICD2, and NICD4.

**Table 3.** Fitted parameters for multiplicative transcription activation model.

| Parameter | Log activation value ($L'_{i,k}$)<sup>a</sup> | Activation value ($m'_{i,k}$)<sup>a</sup> |
| --- | --- | --- |
| NICD1 RAM | 0.326 +/- 0.168 | 1.385 ± 0.233 |
| NICD1 ANK | 0.332 +/- 0.143 | 1.394 ± 0.199 |
| NICD1 CTR | 0.516 +/- 0.186 | 1.675 ± 0.312 |
| NICD2 RAM | -0.026 +/- 0.196 | 0.974 ± 0.191 |
| NICD2 ANK | -0.119 +/- 0.168 | 0.888 ± 0.149 |
| NICD2 CTR | 0.128 +/- 0.196 | 1.136 ± 0.223 |
| NICD3 | 5.027 +/- 0.132 | 152.475 ± 20.127 |
| NICD4 RAM | 0.084 +/- 0.236 | 1.088 ± 0.257 |
| NICD4 ANK | 0.313 +/- 0.156 | 1.368 ± 0.213 |
| NICD4 CTR | -0.516 +/- 0.262 | 0.597 ± 0.156 |

<sup>a</sup>Uncertainties are standard errors on fitted parameters from the covariance matrix.

## Discussion

### RAM and ANK identities determine the DG°RA and DG°MAML, respectively

The ranking of free energies of RAMANK binding to CSL among the four Notch paralogues differ from those of MAML binding to RAMANK:CSL. Notch1 has the most favorable binding free energy for both reactions, whereas the rankings of the other paralogs differ (Notch2 > Notch4 ≥ Notch3 for RAMANK binding, Notch3 ≈ Notch4 > Notch2 for MAML binding). Comparing the free energies of these two binding reactions for RAMANK paralogues with those for RAMANK chimeras indicates that the affinity of RAMANK for CSL is determined by the RAM region (Figure 2) wheras the affinity of MAML for RAMANK:CSL is determined by the ANK domain (Figure 3). This partitioning helps resolve an important question in Notch ternary complex formation: whether or not the ANK domain in the RAMANK:CSL complex is engaged with CSL, as it is in crystal structures of Notch ternary complexes^14,15,18^, or whether it remains dissociated until MAML binds. The dependence of ΔG° for binding of MAML to RAMANK:CSL, and the independence for binding of RAMANK to CSL provides clear thermodynamic evidence that the ANK domain remains dissociated from CSL in the RAMANK:CSL complex, requiring MAML binding to form its interface with CSL.

### Paralogue-specific variations in DG°RA and DG°MAML values and potential sequence origins

The free energies for RAMANK binding to CSL and for MAML binding to RAMANK:CSL vary among Notch paralogues and chimeras. Both *ΔG°_RA_*, and *ΔG°_MAML_* vary over about 1.7 kcal mol^−1^ (Figure 2, 3; Table 1, 2). Under conditions of half-saturation, where occupancy is maximally sensitive to affinity, this range can shift occupancy from 15 to 71 percent. These differences necessarily have their origins in the sequence differences among the four Notch RAMANK regoins. The average pairwise identity among the four ANK paralogues is 62 percent, that among the four RAM regions is 35 percent.

To date, little is known about the specific interactions that stabilize the ANK-CSL and ANK-MAML interfaces, but there have been several studies characterizing the sequence determinants of the RAM-CSL interface, highlighting the important contribution to the affinity of the conserved φWφP motif at the N-terminus of RAM^14,24,25^. Although there is some sequence variation among the φWφP motifs of the four paralogs, these sequence differences do not explain differences in the free energies of the RAMANK:CSL binary complexes. For example, Notch1 and 3 have identical φWφP sequences (LWFP), yet the ΔG°_RA_ values for Notch1 and 3 differ significantly, with Notch3 forming the least stable RAMANK:CSL complex and Notch1 forming the most stable complex. One possible explanation for the decreased stability of the Notch3 RAMANK:CSL complex is substitution of glycine for serine at residue 1672 (Supplemental Figure 4), which has been shown to contribute to the affinity of the RAM peptide with the BTD of CSL^25^. However, given the high conservation around the φWφP motifs of the human Notch1-4 paralogues, it is likely that differences in binary complex stabilities measured here result from sequence differences outside the φWφP motifs.

One sequence feature that has been shown to influence the transcriptional activation by NICD is charge patterning^22^, which can be represented by the parameter κ^26^. Charge patterning has been shown to influence the compaction of RAM and its binding to CSL. The RAM region of Notch3 has the lowest κ value of all the paralogs, which would promote a more expanded structure in this region. This may lower the affinity of the ANK domain for its site on CSL by decreasing the effective concentration of ANK in the unbound state, and may explain why the R3A1 RAMANK:CSL:MAML complex is destabilized by almost 1 kcal mol^−1^ compared to the other constructs containing the N1 ANK domain (Figure 3). However, the expected reciprocal stabilization of the R1A3 RAMANK:CSL:MAML complex relative to R3A3 is not seen.

### The free energy of formation of the Notch RAMANK:CSL:MAML ternary complex

In this study, we have restricted our ITC measurements to reactions with heterodimer stoichiometries. However, the fact that we can separate binding of RAMANK and MAML to CSL provides a method for calculating the free energy of formation of the Notch ternary complex, ΔG°_TC_:

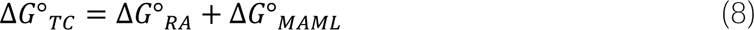

where ΔG°_RA_ and ΔG°_MAML_ are the free energies of binding of RAMANK to CSL (Table 1) and of MAML to RAMANK:CSL (Table 2), respectively. Because the Notch1 ΔG°_RA_ and ΔG°_MAML_ are the most favorable (negative) of all the paralogous values, the Notch1 ΔG°_TC_ is also the most favorable, with a value of −22.4 kcal mol^−1^. For the other three paralogues, the opposite ranking of the ΔG°_RA_ and ΔG°_MAML_ (Notch2 < Notch4 < Notch3 for ΔG°_RA_; Notch3 < Notch4 < Notch2 for ΔG°_MAML_), results in nearly identical ΔG°_TC_ values for Notch2, 3, and 4 (−20.72, −20.82, and −20.70 kcal mol^−1^). That is, Notch1 forms the most stable ternary complex (by about 1.7 kcal mol^−1^), whereas Notch2, 3, and 4 forms ternary complexes of identical (but lower) stability, even though the contributions of RAM and ANK to ternary complex stability differ. It is possible that the differences in the RAM and ANK affinities give rise to paralogue specific differences in signaling outcomes.

### The relationship between transcriptional activation and the stability of RAMANK:CSL:MAML ternary complexes

The transcription level at Notch-responsive promoters should be proportional to the amount of Notch ternary complex assembly, which in turn, should be proportional to the ΔG°_TC_, especially at subsaturating levels of NICD. Our ITC and transcriptional activation data provides an opportunity to quantitatively examine this relationship. Values for the four Notch paralogues support such a correlation: Notch1 shows roughly three times the transcriptional activation as Notch2, 3, and 4, which are all roughly the same (Figure 4); as described in the preceeding section, values of ΔG°_TC_, show the same distribution, with the Notch1 paralog forming the most stable ternary complex, and Notch 2, 3, and 4 having similar decreased stabilities.

Transcriptional activation from the ten chimeric NICD constructs vary considerably; we measured ΔG°_RA_ and ΔG°_MAML_ values for six of these constructs, allowing us to further test the relationship between transcriptional activation and ternary complex stability (Figure 6A). Although the ΔG°_TC_, values for the paralogous and chimeric RAMANK constructs do show a positive correlation with transcriptional activation values, there are large deviations from a linear fit. These deviations may result from variations in contributions to transcription activation from the four CTRs, which are considerable (Figure 5A) and are not accounted for in the ITC data. Indeed, all constructs containing the NICD1 CTR have an observed activation above the regression line, consistent with the large CTR1 coefficient in our multiplicative activation model (Figure 5). Likewise, the R4A1C4 chimera has an activation below the regression line, consistent with the small CTR4 coefficient in our model.

**Figure 6.**
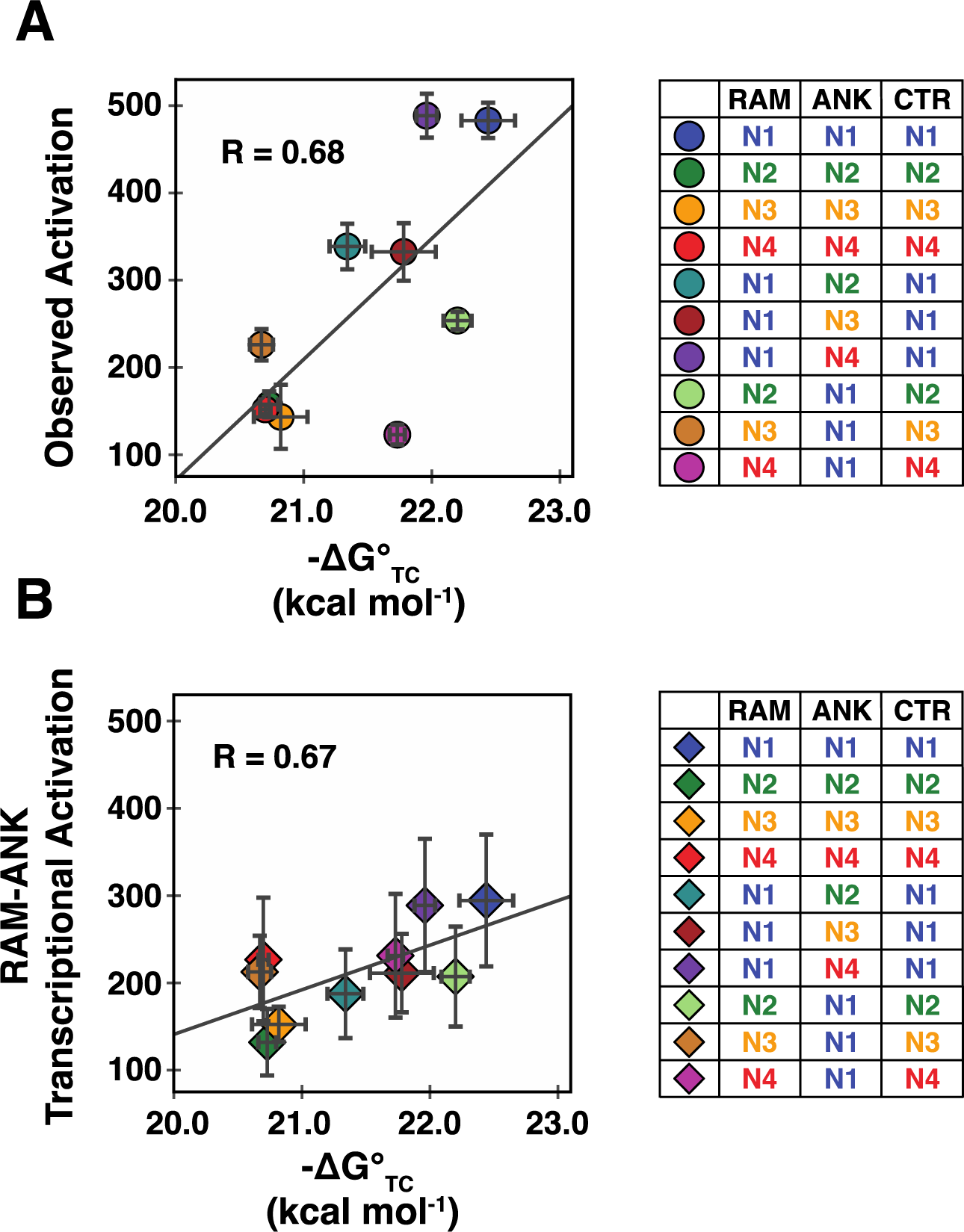
Correlation of free energy values for RAMANK:CSL:MAML ternary complex formation with transcriptional activation. ΔG°_TC_ values were calculated by summing ΔG°_RA_ and ΔG°_MAML_ values (equation 8). A) ΔG°_TC_ values versus measured transcriptional activation for NICD paralogues and chimeras. B) ΔG°_TC_ values versus the component of the transcriptional activation that is attributable to the RAM and ANK regions, based on our model parameters. R-values are Pearson correlation coefficients.

To attempt to control for varations in transcriptional activation from the CTRs, we calculated activation for the ten constructs for which we collected ITC data using only the coefficients for the RAM and ANK regions from our model, along with the intercept coefficient for NICD3. These partial activation values, which can be viewed as predicted activations for constructs with variable RAM and ANK regions but the same (Notch3) CTR, indeed show a positive correlation with ΔG°_TC_ values (Figure 6B). Although the Pearson correlation coefficient is the same as for the full (i.e., measured) transcriptional activation values (Figure 6A), the deviations from the regression line are much smaller for the partial activations (41 units for partial activation values, 106 for the full values). Whereas deviations for the full transcriptional activation parameters are significantly larger than than the uncertainties in activation (Figure 6A), deviations for the partial activations are within their uncertainties. On the whole, the relationship in Figure 6B supports the hypothesis that transcriptional activation is determined by ΔG°_TC_, with a correlation that is is probably as good as can be expected given the large errors on the partial activation values.

### Similarities and differences between binding free energies and transcriptional activation contributions of the RAM and ANK regions

As described above, ternary complex stabilities are positively correlated with transcriptional activation values. Since ternary complex stabilities are obtained by summing the ΔG°_RA_ and ΔG°_MAML_ values, and since these binding energies appear to be determined by RAM and ANK binding respectively, these ΔG° values should correlate with the fitted RAM and ANK activation coefficients from the multiplicative model. For the RAM region, there are indeed similar trends between the RAM activation parameters and ΔG°_RA_ values. Notch1 has the largest fitted RAM activation parameter, and constructs with Notch1 RAM show the most favorable ΔG°_RA_ values. Likewise, Notch 3 and 4 have similar activation parameters, and constructs with Notch 3 and 4 RAM regions have similar ΔG°_RA_ values. However, Notch2 provides an exception to this correlation: the Notch2 RAM activation parameter is below (albeit only slightly) those of Notch3 and Notch4, yet constructs with Notch2 RAM have more favorable ΔG°_RA_ values than do Notch 3 and Notch4.

There are also similarities between ANK activation parameters and ΔG°_MAML_. Notch1 has the largest fitted ANK activation parameter, and constructs with Notch1 ANK show the most favorable ΔG°_MAML_ values. Notch2 has the smallest fitted ANK activation parameter, and constructs with Notch2 ANK show the least favorable ΔG°_MAML_ values. However, Notch4 provides an exception to this correlation: the Notch4 ANK activation parameter is below (albeit only slightly) those of Notch3 and Notch2, yet constructs with Notch4 ANK have more favorable ΔG°_RA_ values than do Notch 2 and Notch3.

For both the RAM and ANK regions, the correlation between binding energies and activation parameters are consistent with these two interactions being important for ternary complex assembly leading to transcriptional activation. However the origins of the noted discrepancies are unclear. It is possible that other proteins interact differentially with these domains to modulate transcriptional activation. For the RAM regions, co-repressor proteins may serve this role, as corepressors are thought to be displaced by the Notch RAM regions.

### Contribution of CTRs to transcriptional activation

As described above, the largest variations in fitted coefficients for transcriptional activation are for the CTRs (Figure 5A). Notch1 has the highest CTR coefficient, Notch2 and 3 have intermediate values, and Notch4 has a low coefficient. This ranking is consistent with previous studies that have examined the sequence features and relative contributions of CTRs to Notch signaling. Notch1 has been reported to have a potent TAD in its CTR^16^, whereas Notch4 lacks a TAD in its CTR. NICD2’s CTR has previously been shown to be interchangeable with that of Notch1^27^; in part, this is consistent with the observation here that the Notch2 CTR activation coefficient is second-largest among the four, although it is significantly smaller than the Notch1 CTR, and is rather close to that of Notch3. Though Notch3 has a TAD in its CTR, it has been described as non-conventional^17^, which may be related to the lower ranking of the Notch3 CTR activation coefficient compared with Notch1.

In conclusion, our analysis of the thermodynamics of paralog-specific NTC formation and the role of individual regions in determining the strength of signaling is a significant step forward in our understanding of the mechanisms behind paralog-specific Notch signaling. This work demonstrates that the thermodynamics of NTC formation does influence signaling in a paralog-specific way, but that the CTRs also play a major role in activation. Further studies that include the role of the NICD CTRs are warranted.

## Author Contributions

KMR prepared recombinant protein samples and designed ITC experiments, fitted isotherms, and analyzed and interpreted data. KMR designed and performed transcriptional activation assays and analyzed and interpreted data. KMR and DB developed the independent multiplicative model for analyzing transcriptional activation data. KMR fitted the transcriptional activation data to the independent multiplicative model. KMR and DB analyzed and interpreted data from the independent multiplicative model. KMR and DB prepared figures and tables. KMR and DB wrote, read, and edited the manuscript.

## Materials and Methods

### Notch RAMANK and CSL expression and purification

The RAMANK regions of the four human Notch paralogues (Table S3) were subcloned into the pMAL-c2x expression vector using HiFi DNA Assembly (New England BioLabs). Constructs were expressed in BL21(DE3) *E. coli* cells as N-terminal MBP-fusions with N-terminal His_12_ tags. Cells were grown at 37 °C in LB containing 0.1 g/L ampicillin to an OD_600_ of 0.8. Isopropyl-β-D-1-thiogalactopyranoside (IPTG) was added to 1 mM, the temperature was reduced to 20 °C, and cells were grown overnight.

Cells were centrifuged at 5000xg and pellets were resuspended in Cell Resuspension Buffer (25 mM Tris pH 8.0, 50 mM NaCl, 0.5 mM TCEP). Cells were disrupted via sonication (2s on, 2s off, 5min) on ice and cell debris was pelleted by centrifugation at 15,000xg for 30min at 4 °C. Clarified lysates were brought to 2 mM MgCl_2_ and DNaseI and benzonase (1 mg/ml, 500 units).

Following a one-hour incubation at room temperature, lysates were brought to 250 mM NaCl and applied to a benchtop column containing Ni-NTA resin (Qiagen) equilibrated in Equilibration Buffer (25 mM Tris pH 8.0, 250 mM NaCl, 0.5 mM TCEP), washed with Equilibration Buffer containing 20mM imidazole, and eluted with Equilibration Buffer containing 250mM imidazole. Eluted MBP-fusion proteins were cleaved with TEV protease during dialysis into TEV Cleavage Buffer (25mM sodium phosphate pH 7.0, 100 mM NaCl, 0.5 mM TCEP) overnight at 4 °C. Cleaved proteins were reapplied to Ni-NTA resin equilibrated in TEV Dialysis Buffer to remove cleaved His_12_-MBP and any uncleaved protein. The flow-through was loaded onto a HiTrapQ HF anion exchange column (Cytiva) and eluted with a linear gradient from 0 to 1M NaCl. Pure protein fractions were pooled and dialyzed overnight at 4 °C into Storage Dialysis Buffer (25 mM sodium phosphate pH 7.0, 150 mM NaCl, 0.5 mM TCEP). Human CSL spanning residues 25-449 was expressed and purified as described for the Notch RAMANK constructs, substituting the final anion exchange column with a HiTrapSP HF cation exchange column (Cytiva). All purified (RAMANK constructs, CSL, and MAML, see below) were either used immediately or flash frozen in 1mL aliquots using liquid nitrogen.

### MAML1 expression and purification

Human MAML1 spanning residues 8-76 was cloned into a pET-SUMO vector and a C-terminal tryptophan was added for concentration determination. Protein was expressed using BL21(DE3) cells at 37 °C with LB containing 50 µg/mL kanamycin at 37°C until OD_600_ was ∼0.8 and induced with IPTG as described for RAMANK constructs. Cells were centrifuged at 5000xg for 10min. Pellets were resuspended, sonicated, and centrifuged as described above. Lysate pellets, which contain the majority of SUMO-MAML1W, were subjected to two washes in 50 mL of 100 mM Tris pH 7.4, 5 mM EDTA, 0.5 mM TCEP, 1% Triton X-100, 1 M urea followed by two washes in 50 mL of 100 mM Tris pH 7.4, 0.5 mM TCEP. In each wash, SUMO-MAML1W pellets were dispersed using a dounce homogenizer and re-pelletted at 15000xg for 30 minutes. The final pellet was resuspended with 8 M urea, 25 mM Tris pH 8.0, 250 mM NaCl, 0.5 mM TCEP and batch-bound to Ni-NTA resin equilibrated with the same buffer overnight at 4 °C with gentle rotation.

After overnight binding, the resin mixture was transferred to a column and was washed with the same resuspension buffer, and SUMO-MAML1W was refolded on the resin by washing with Refold Buffer (25 mM Tris pH 8.0, 250 mM NaCl, 0.5 mM TCEP). Protein was then eluted with Refold Buffer containing 250 mM imidazole, and EDTA (pH 8.0) was immediately added to a concentration of 0.1 M. SUMO protease Ulp1 was added to the eluted protein and dialyzed at 4 °C in 50 mM Tris pH 8.0, 150 mM NaCl, 0.5 mM TCEP 5% glycerol for 8 hours, and subsequently dialysed into 25 mM sodium phosphate pH 8.0, 50 mM NaCl, 0.5 mM TCEP overnight at 4 °C.

Following dialysis, insoluble aggregates were removed by centrifugation at 18,000xg for 30min. Clarified dialysate was loaded onto a cation exchange column (HiTrapSP HF, Cytiva) and eluted with a linear gradient from 0-1 M NaCl in 25 mM sodium phosphate pH 8.0 and 0.5 mM TCEP. Fractions containing pure MAML1W were pooled and dialyzed at 4 °C overnight into Storage Dialysis Buffer.

### Isothermal Titration Calorimetry

Proteins were dialyzed overnight at 4 °C into 25 mM sodium phosphate pH 7.0, 150 mM NaCl, 0.5 mM TCEP. After dialysis, proteins were filtered using 0.22 µm PVDF membranes and concentrations were determined using UV absorbance and extinction coefficients obtained from the amino acid sequence using the ExPASy ProtParam tool.

ITC thermograms were obtained at 25 °C using a MicroCal VP-ITC instrument. To monitor binary complex formation, 20 μM RAMANK was titrated into 2 μM CSL using 8 μL injections at 5min intervals. For MAML1 binding to RAMANK:CSL complexes, 20 μM MAML was titrated into 2 μM CSL and 20 μM RAMANK using 8 μL injections at 5 min intervals. Heat peaks from the raw thermogram were integrated using NITPIC^30^ and the integrated thermogram was fitted with a single-site binding model using SEDPHAT^31^. Fitted parameters are reported as an average of at least three experiments; reported uncertainties are standard error of the mean.

### Transcriptional activation assays

NICD open reading frames (NICD1, 1758-2555; NICD2, 1699-2471; NICD3, 1663-2321; and NICD4, 1472-2003) were cloned into the pcDNA3.1(+) mammalian expression vector with an added C-terminal FLAG tag. Chimeric NICDs were created from native NICDs using HiFi DNA Assembly (New England Biolabs). Boundaries for chimeric constructs are given in Table S3.

HeLa cells were maintained in Dulbecco’s modified Eagle’s medium (Gibco) with 10% (v/v) heat-inactivated fetal bovine serum (FBS, Corning) and 1% (v/v) penicillin/streptomycin (Gibco) at 37 °C and 5% CO_2_ in a humidified incubator. 24 hours prior to transfection, cells were seeded into 12-well plates at a density of 4 x 10^5^ cells per well and were incubated at 37 °C. Cells were transiently transfected with 225 ng TP1-luc reporter plasmid, 75 ng Renilla transfection control plasmid, and 200 ng NICD-expressing plasmid (or empty pcDNA3.1(+) negative control plasmid) using Lipofectamine LTX with PLUS reagent (Life Technologies). Cells were lysed 44-48 hours post-transfection by adding 250 μL Passive Lysis Buffer (Promega) to each well. Luciferase activity of lysates was measured using the Dual Luciferase Assay Reagents (Promega) on a GloMax Multi microplate luminometer (Promega). Firefly luciferase activities for NICD constructs and negative control were corrected for variations in transfection efficiency by dividing by Renilla luciferase activities. These ratiometric activities were normalized by the Renilla-corrected negative control. Luciferase activities for each construct were measured in triplicate (in three separate wells) for each transfection, and transfections were repeated at least three times. Reported transcriptional activation values are the average of the three or more transfections performed in triplicate; reported uncertainties are standard error of the mean.

Expression of each NICD was confirmed with Western blotting. NICD-FLAG-containing lysates were separated on a 7.5% acrylamide Mini-PROTEAN TGX gels (Bio-Rad) and transferred to PVDF membranes using a Trans-Blot Turbo semi-dry transfer system (Bio-Rad). Blots were blocked with 5% BSA in PBS-T (PBS containing 0.05% Tween-20) at room temperature with gentle agitation for four hours. Membranes were probed with primary antibodies overnight at 4 °C in blocking buffer with gentle agitation. Anti-FLAG primary antibody (Sigma F7425) was diluted 3000-fold; anti-GAPDH primary antibody (Cell Signaling 14C10) was diluted 5000-fold. Following incubation with primary antibodies, membranes were washed at least six times with PBS-T. Blots were probed with anti-rabbit IgG1-HRP secondary antibody (Cell Signaling 7074) at 1:10000 dilution in blocking buffer for 1 hour at room temperature with gentle agitation and were then washed six times with PBS-T. Bands were visualized using ECL Prime Western Blotting Detection Reagents (Amersham) and imaged on autoradiography film.

## Key Resources Table

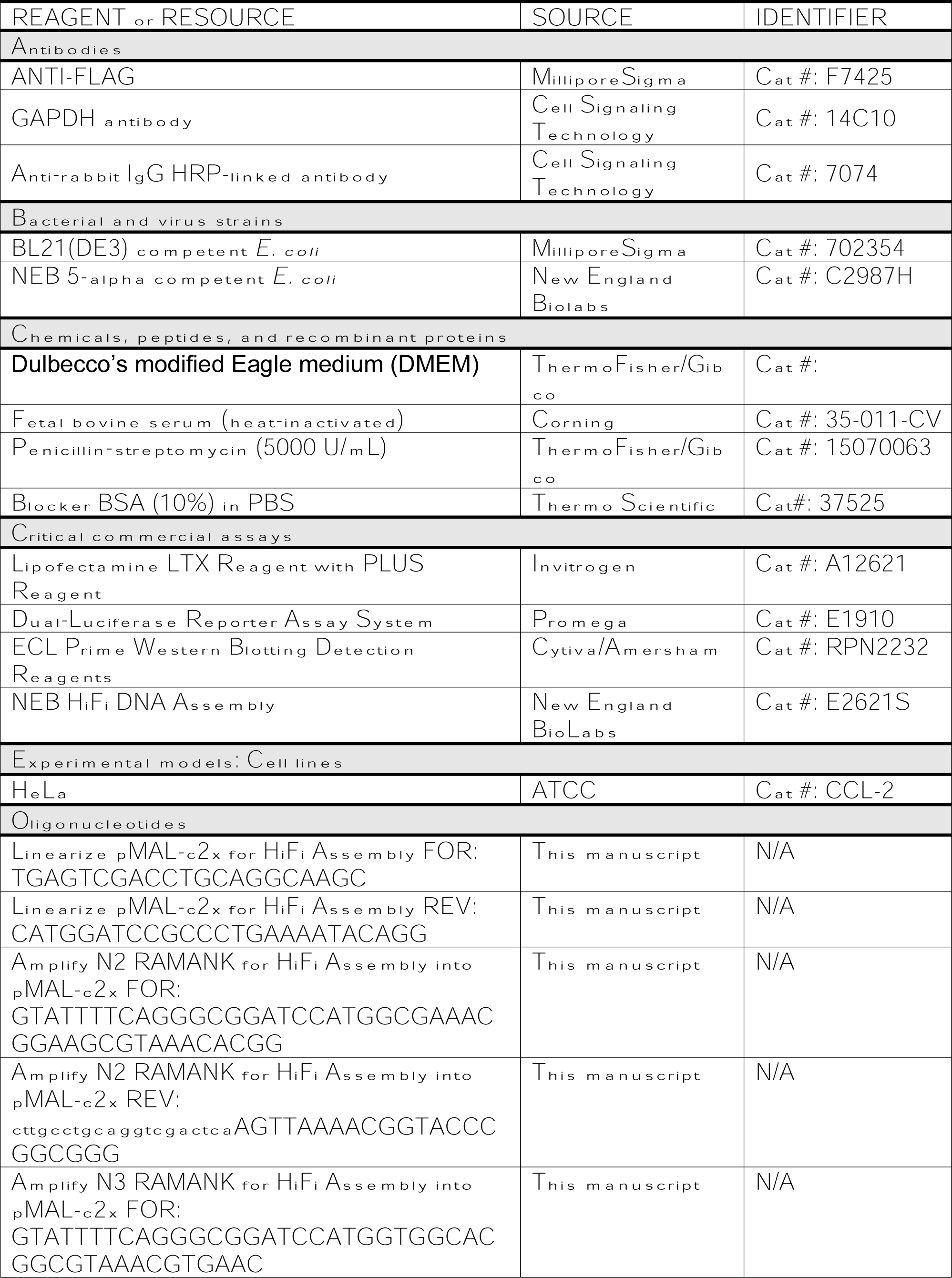
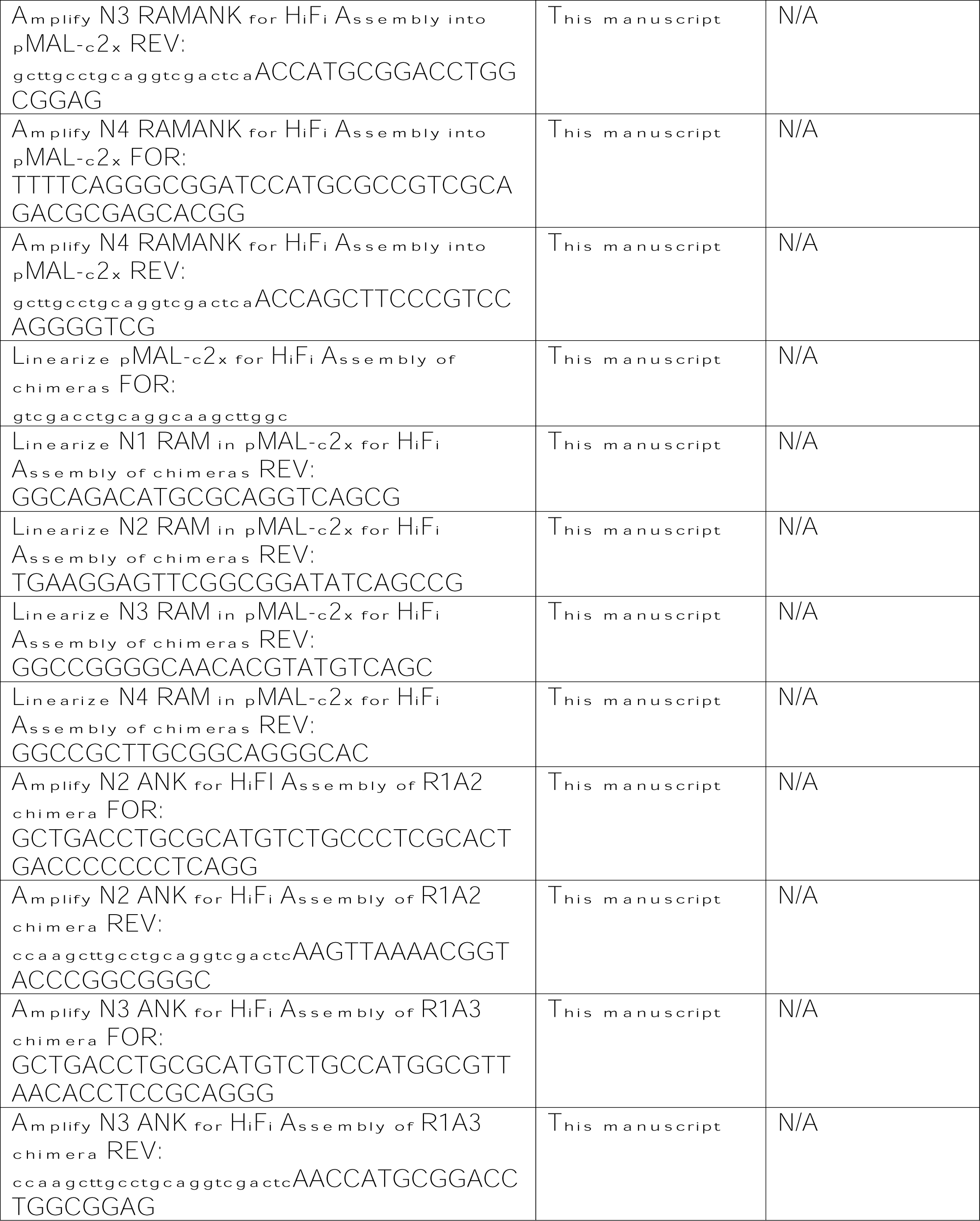
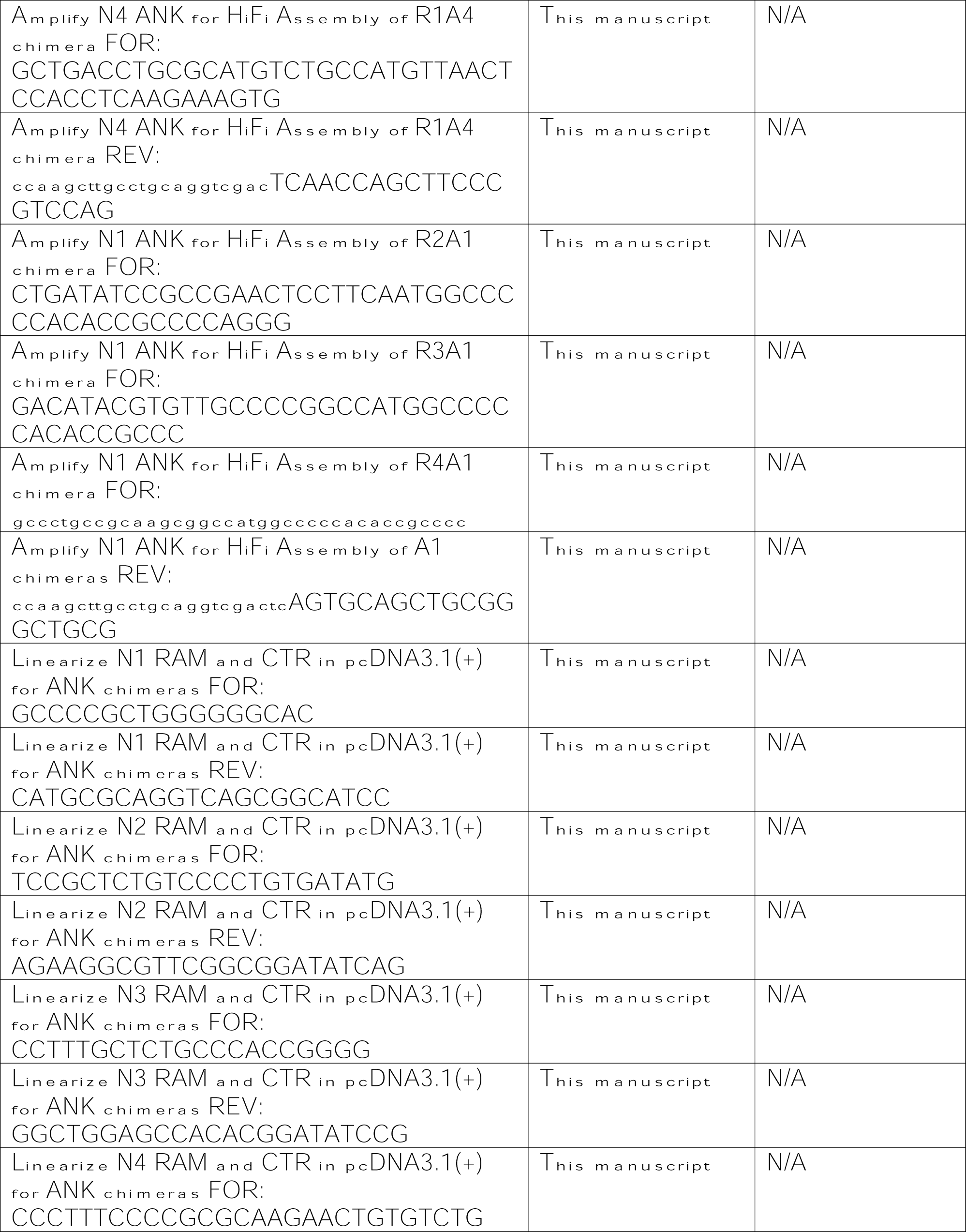
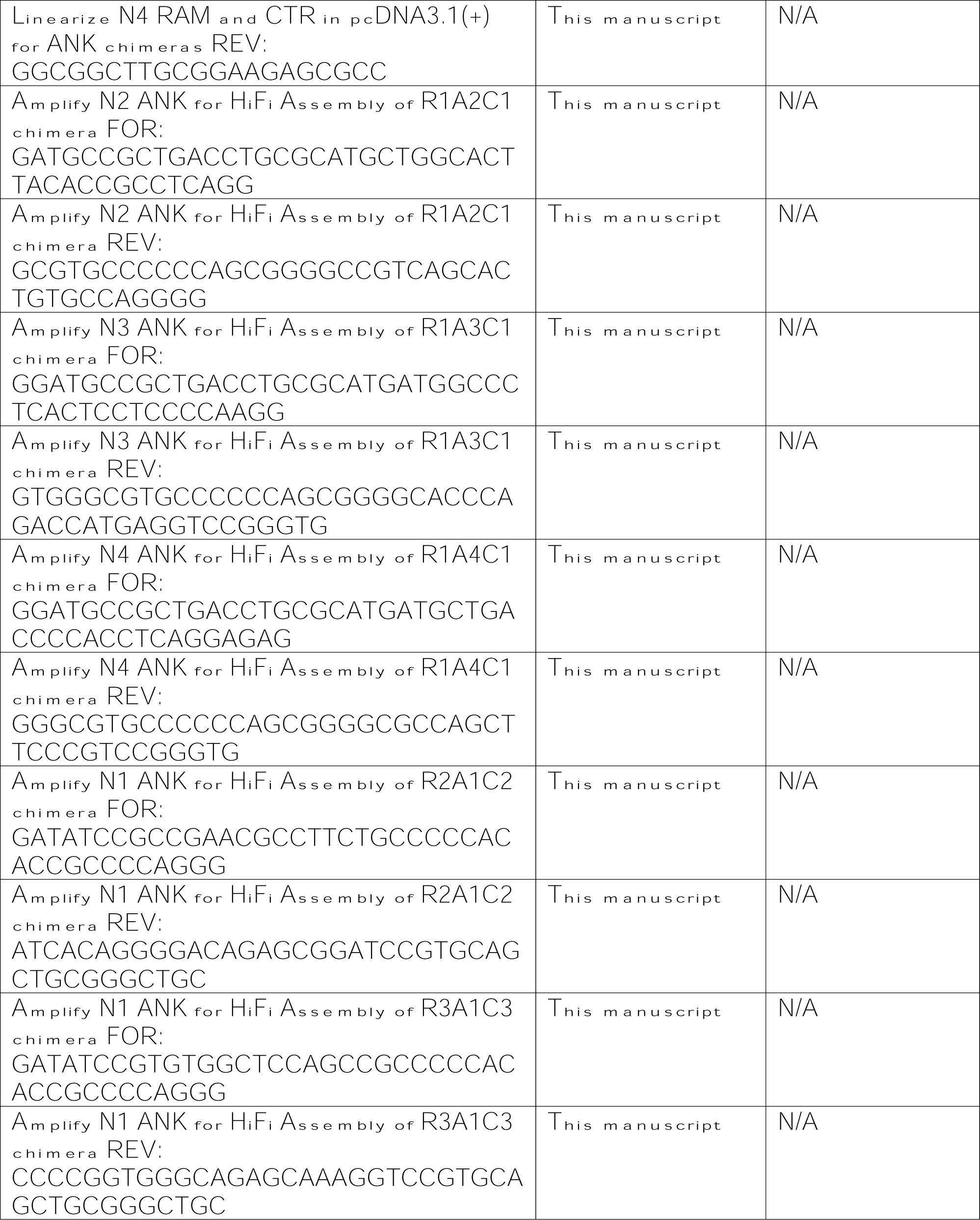
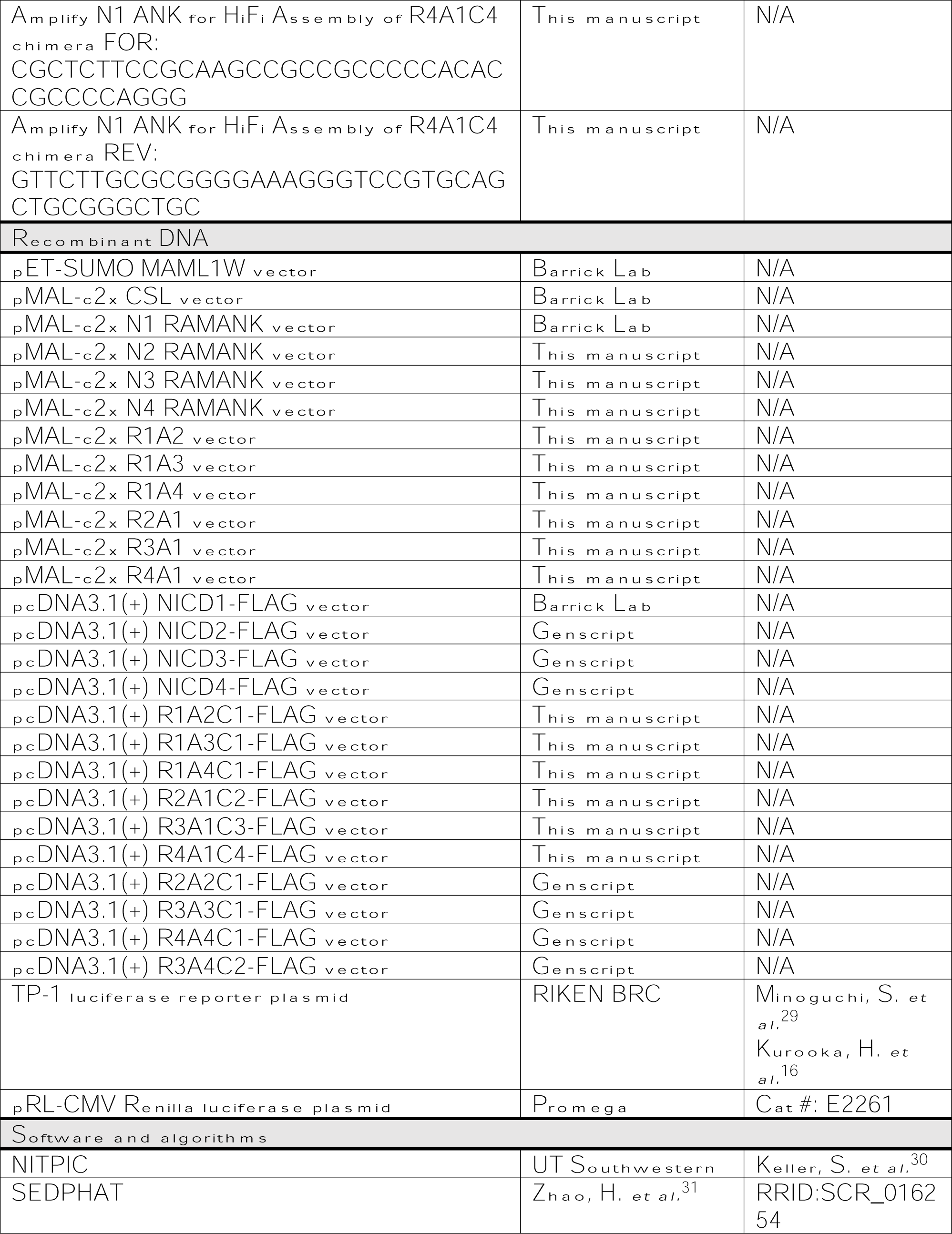
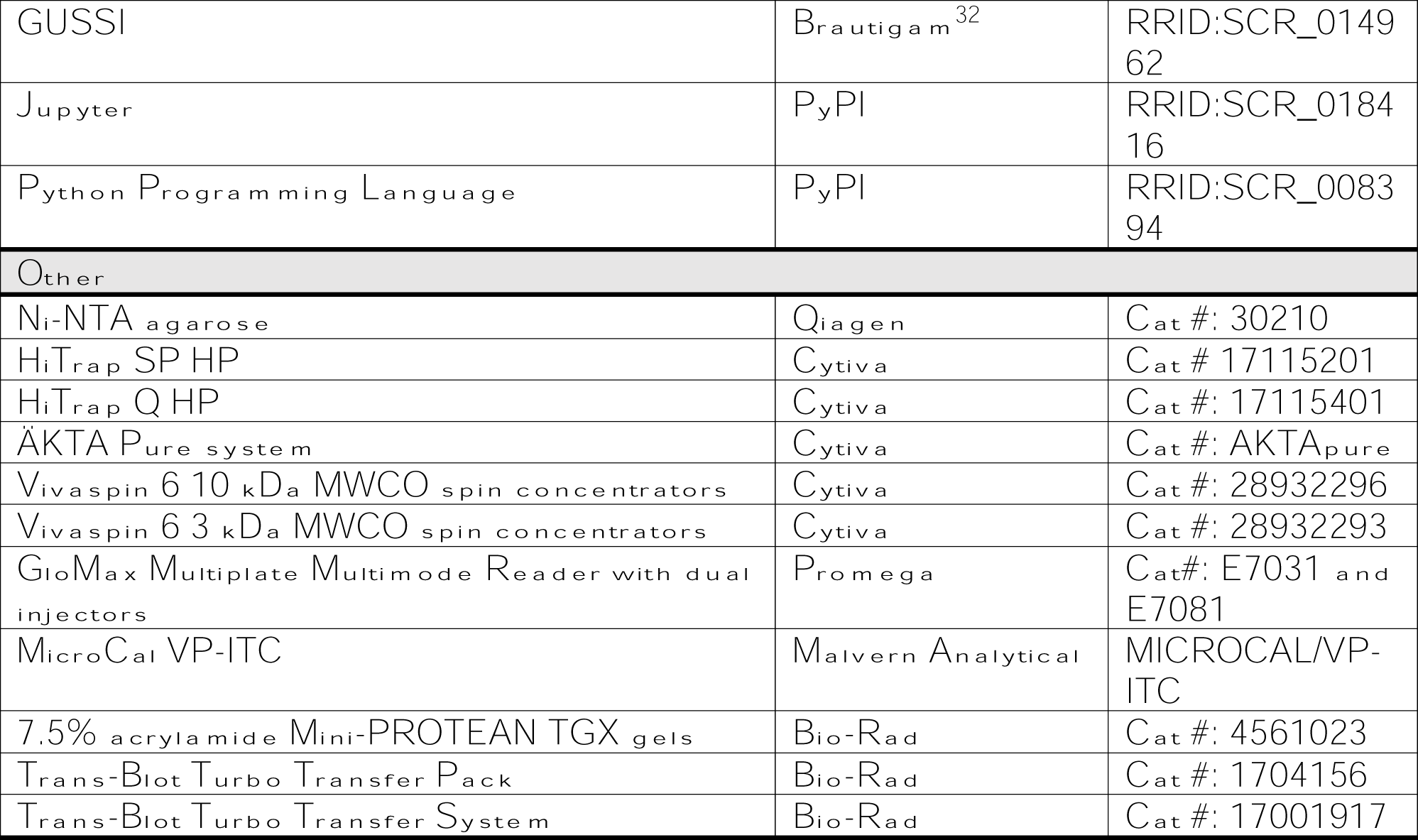

## Footnotes

1 The 14-by-12 data matrix does not have full column rank; this would be true even if we were to collect data for all 4^3^=64 possible chimeric constructs.

