## Supplementary figures and images for "Unravelling Paralog-Specific Notch Signaling through Ternary Complex Stability and Transcriptional Activation Measurements Using Chimeric Receptors"

### Supplemental Figure 1

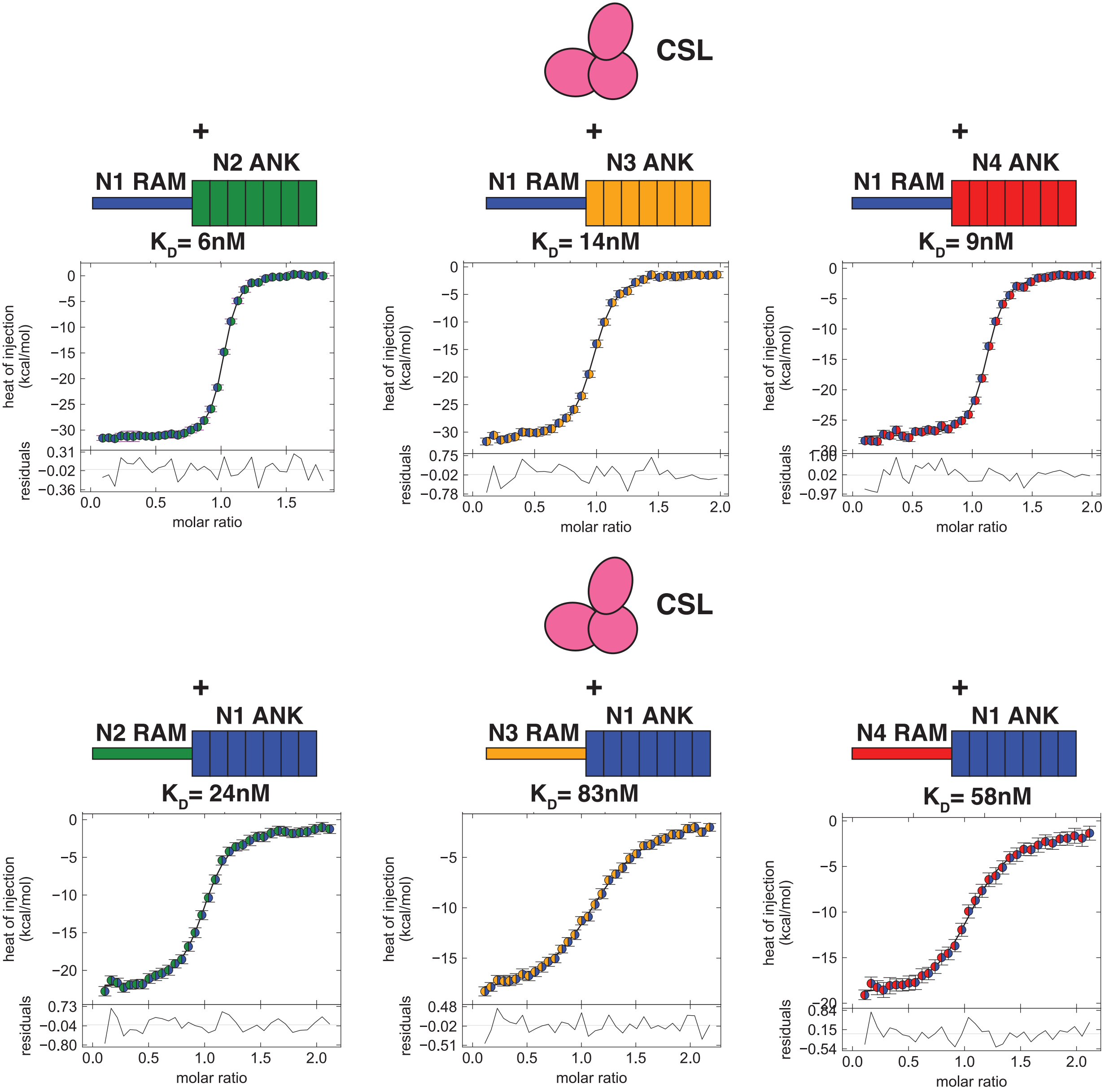

### Supplemental Figure 2

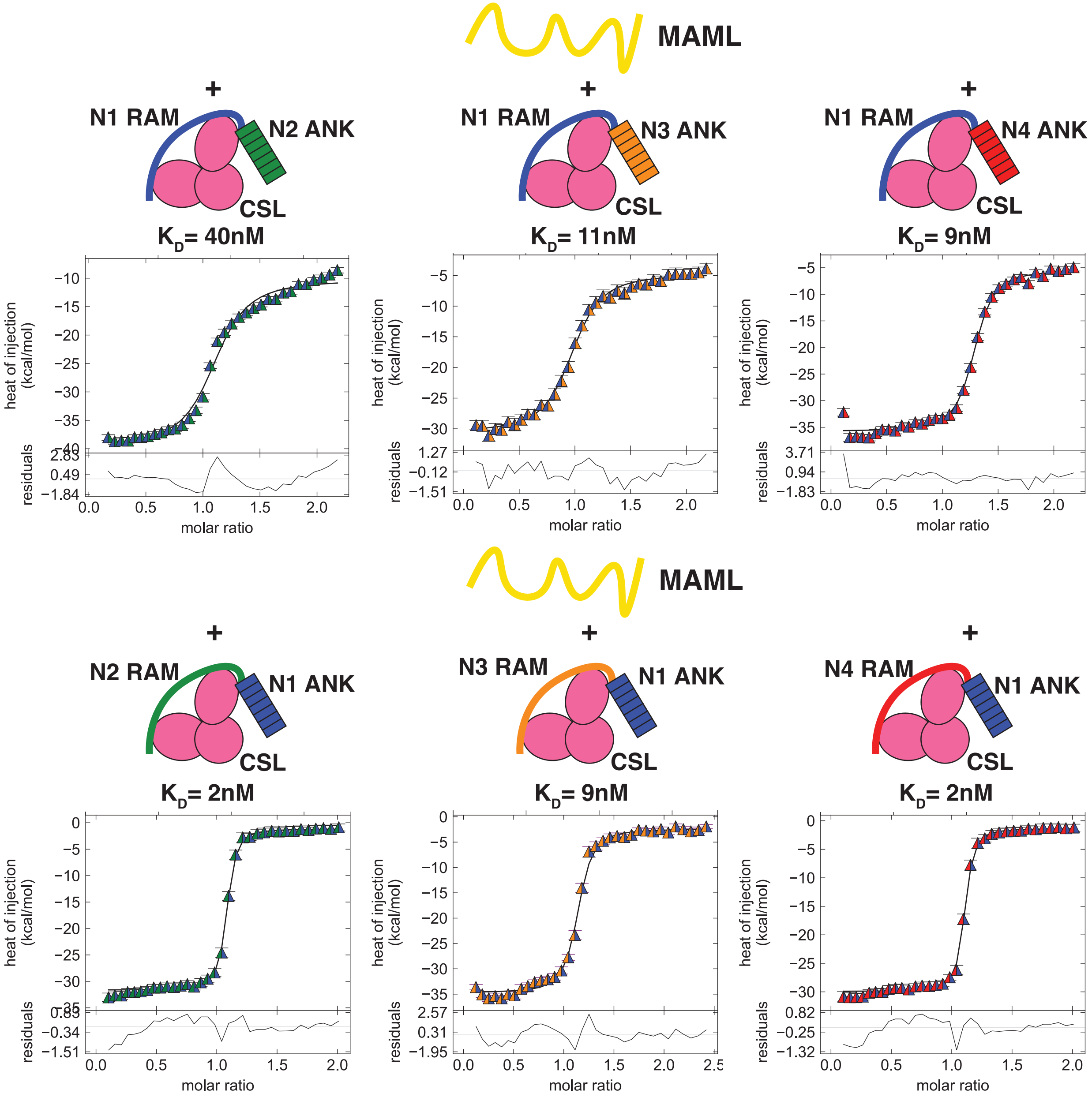

### Supplemental Figure 3

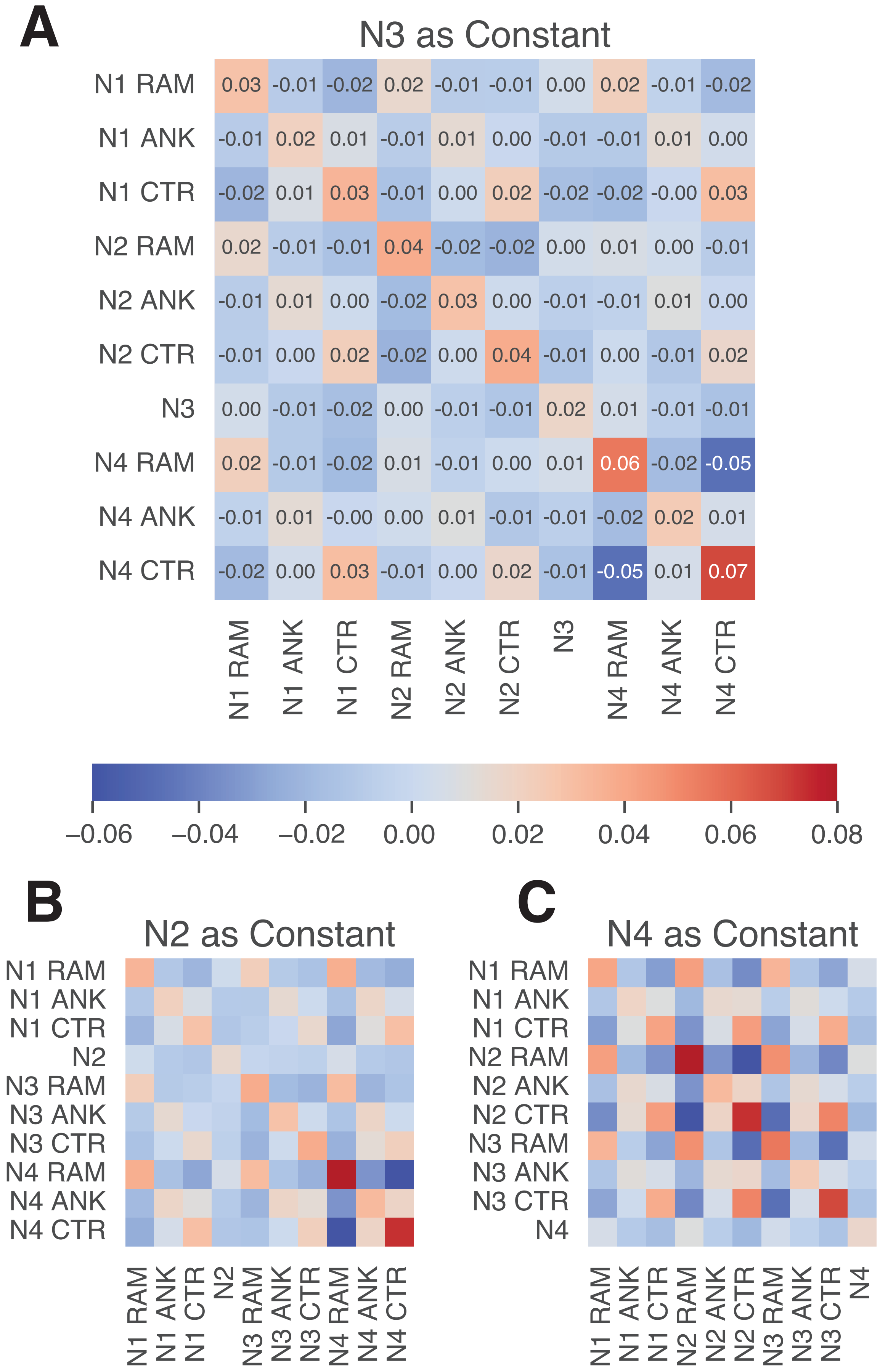

### Supplemental Figure 5

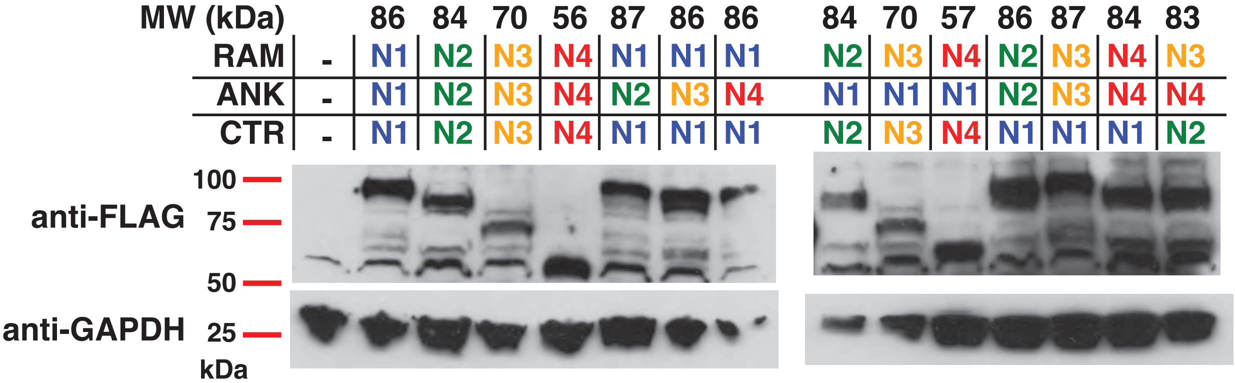
