## Supplemental Figure 4 for "Unravelling Paralog-Specific Notch Signaling through Ternary Complex Stability and Transcriptional Activation Measurements Using Chimeric Receptors"

ΦWΦP RAM

```
N1  1758 ..RKRRRQHGQLWFPEGFKVSEA...SKKKRREPLGEDSVGLKPLKNASD....GALMDD 1808
N2  1699 .MAKRKRKHGSLWLPEGFTLRRDAS..NHKRREPVGQDAVGLKNLSVQVSE..ANLIGTG 1753
N3  1663 MVARRKREHSTLWFPEGFSLHKDVASGHKGRREPVGQDALGMKNMAKGESL......MGE 1716
N4  1472 ..RRRREHGALWLPPGFTRRPRTQSAPHRRRPPLGEDSIGLKALKPKAEVDEDGVVMCS 1529
```

RAM

```
N1  1809 NQNEWGDE.DLETKKFRFEEPVVLPDLDDQTDHRQWTQQHLDAADLRM.SAMAPTPPQGE 1866
N2  1754 TSEHWVDDEGPQPKKVKAEDEALLSEEDDPIDRRPWTQQHLEAADIRRTPSLALTPPQAE 1813
N3  1717 VATDWMDTECPEAKRLKVEEPGMGA..EEAVDCRQWTQHHLVAADIRVAPAMALTPPQGD 1774
N4  1530 G.........PE....EGEEVGQAEETGPPSTCQLWSLSGGC....GALPQAAMLTPPQE 1572
```

Disordered AR1

```
N1  1867 VDADCMDVNVRGPDGFTPLMIASCSGGGLETGNSEE...EEDAPAVISDFIYQGASLHNQ 1923
N2  1814 QEVDVLDVNVRGPDGCTPLMLASLRGGSSDL.SDEDEDAEDSSANIITDLVYQGASLQAQ 1872
N3  1775 ADADGMDVNVRGPDGFTPLMLASFCGGALEPMPTEEDEADDTSASIISDLICQGAQLGAR 1834
N4  1573 SEMEAPDLDTRGPDGVTPLMSAVCCGEVQSGTF...QGAWLGCPEPWEPLLDGGACPQAH 1629
```

AR2 AR3

```
N1  1924 TDRTGETALHLAARYSRSDAAKRLLEASADANIQDNMGRTPLHAAVSADAQGVFQILIRN 1983
N2  1873 TDRTGEMALHLAARYSRADAAKRLLDAGADANAQDNMGRCPLHAAVAADAQGVFQILIRN 1932
N3  1835 TDRTGETALHLAARYARADAAKRLLDAGADTNAQDHSGRTPLHTAVTADAQGVFQILIRN 1894
N4  1630 TVGTGETPLHLAARFSRPTAARRLLEAGANPNQPDRAGRTPLHAAVAADAREVCQLLLRS 1689
```

AR3 AR4 AR5

```
N1  1984 RATDLDARMHDGTTPLILAARLAVEGMLEDLINSHADVNAVDDLGKSALHWAAAVNNVDA 2043
N2  1933 RVTDLDARMNDGTTPLILAARLAVEGMVAELINCQADVNAVDDHGKSALHWAAAVNNVEA 1992
N3  1895 RSTDLDARMADGSTALILAARLAVEGMVEELIASHADVNAVDELGKSALHWAAAVNNVEA 1954
N4  1690 RQTAVDARTEDGTTPLMLAARLAVEDLVEELIAAQADVGARDKWGKTALHWAAAVNNARA 1749
```

AR5 AR6 AR7

```
N1  2044 AVVLLKNGANKDMQNNREETPLFLAAREGSYETAKVLLDHFANRDITDHMDRLPRDIAQE 2103
N2  1993 TLLLLKNGANRDMQDNKEETPLFLAAREGSYEAAKILLDHFANRDITDHMDRLPRDVARD 2052
N3  1955 TLALLKNGANKDMQDSKEETPLFLAAREGSYEAAKLLLDHFANREITDHLDRLPRDVAQE 2014
N4  1750 ARSLLQAGADKDAQDNREQTPLFLAAREGAVEVAQLLLGLGAARELRDQAGLAPADVAHQ 1809
```

AR7

```
N1  2104 RMHHDIVRLLDEYNLVRSPQLH....G.. 2126
N2  2053 RMHHDIVRLLDEYNVTPSPPGT....VLT 2077
N3  2015 RLHQDIVRLLDQPSGPRSPPGP....HG. 2038
N4  1810 RNHWDLLTLLEGAGPPEARHKATPGREAG 1838
```
