## Supplemental Table 1 for "Unravelling Paralog-Specific Notch Signaling through Ternary Complex Stability and Transcriptional Activation Measurements Using Chimeric Receptors"

**Supplemental Table 1. Boundaries of RAM, ANK, and CTR regions for the four Notch paralogs used in this study**

| | RAM region | ANK domain | CTR |
|---|---|---|---|
| **NICD1** | 1758-1857 | 1858-2126 | 2127-2555 |
| **NICD2** | 1699-1804 | 1805-2077 | 2078-2471 |
| **NICD3** | 1663-1766 | 1767-2038 | 2039-2321 |
| **NICD4** | 1472-1566 | 1567-1838 | 1839-2003 |
